# The simplicity of protein sequence-function relationships

**DOI:** 10.1101/2023.09.02.556057

**Authors:** Yeonwoo Park, Brian P.H. Metzger, Joseph W. Thornton

## Abstract

How complicated is the genetic architecture of proteins – the set of causal effects by which sequence determines function? High-order epistatic interactions among residues are thought to be pervasive, making a protein’s function difficult to predict or understand from its sequence. Most studies, however, used methods that overestimate epistasis, because they analyze genetic architecture relative to a designated reference sequence – causing measurement noise and small local idiosyncrasies to propagate into pervasive high-order interactions – or have not effectively accounted for global nonlinearity in the sequence-function relationship. Here we present a new reference-free method that jointly estimates global nonlinearity and specific epistatic interactions across a protein’s entire genotype-phenotype map. This method yields a maximally efficient explanation of a protein’s genetic architecture and is more robust than existing methods to measurement noise, partial sampling, and model misspecification. We reanalyze 20 combinatorial mutagenesis experiments from a diverse set of proteins and find that additive and pairwise effects, along with a simple nonlinearity to account for limited dynamic range, explain a median of 96% of total variance in measured phenotypes (and >92% in every case). Only a tiny fraction of genotypes are strongly affected by third- or higher-order epistasis. Genetic architecture is also sparse: the number of terms required to explain the vast majority of variance is smaller than the number of genotypes by many orders of magnitude. The sequence-function relationship in most proteins is therefore far simpler than previously thought, opening the way for new and tractable approaches to characterize it.

## Introduction

If we had complete knowledge of a protein’s genetic architecture – the set of causal rules by which its sequence determines its functions – then we could predict and understand the functional and evolutionary consequences of any variant sequence. Whether such knowledge is possible in practice depends on the extent of epistatic interactions. If all residues in a protein acted independently, knowing the effects of point mutations on any genetic background would suffice to understand the functional contribution of every possible residue and predict the function of every possible sequence; moreover, any mutation’s evolutionary fate would be independent of the genetic context in which it may arise. Such a simple genetic architecture could be reconstructed by moderate-throughput experiments. At the opposite extreme, pervasive high-order epistasis would cause a mutation’s effect to vary idiosyncratically across genetic backgrounds, and the evolutionary fate of any mutation would change unpredictably with each sequence substitution. Assessing the genetic architecture would require exhaustive characterization of all possible sequences.

High-throughput methods for characterizing large libraries of protein variants have made it possible to directly assess the complexity of sequence-function relationship. Studies to date disagree on the extent of epistasis within proteins. Some report extensive high-order interactions^1–7^, while others find that they account for less than 10% of functional variance among sequences^8–18^. Even pairwise interactions are strong and widespread in some studies^7,12,18–22^ but weak and rare in others^9,16,23,24^. Some studies report a sparse genetic architecture in which a small fraction of possible amino acids and interactions dictate the function^13,16^, but others point to a more complex mapping in which many determinants of small effect contribute to function^7,18,20,22^.

These discrepancies may arise from the use of different methods to characterize epistasis. Two aspects of widely used approaches can overestimate the complexity of genetic architecture. First, many studies have analyzed combinatorial mutagenesis data from a reference-based framework, which designates a single sequence as wild-type. If a mutation’s effect when introduced into a variant differs from its effect on the wild-type, the deviation is attributed to epistasis, even though it may reflect propagation of error from measurement noise or local idiosyncrasies in the wild-type architecture^25^. Second, many studies have not fully accounted for global nonlinearities in the relationship between sequence and function^26^. When this nonspecific epistasis is not incorporated, spurious amino acid interactions must be invoked to explain why every mutation’s effect varies across genetic backgrounds^11,27,28^.

Advances have been made in both areas of concern, but current methods have major limitations. Fourier analysis^29,30^, also known as simplex encoding^31^ or graph Fourier transform^32^, is reference-free: instead of focusing on the effects of states on a particular sequence, it captures their average effects across sequence space. But the application of Fourier analysis has been mostly limited to datasets that sample just two states per site, because the multi-state formalism is complicated and has no straightforward interpretation. For example, when all 20 amino acids are present, they must be recoded into 19 Fourier coefficients using Hadamard matrices or graph Fourier bases, and the resulting model terms do not correspond to any genetically or biochemically meaningful quantities. Another formalism, background-averaging^2,25,33,34^, is a modified reference-based analysis in which the effects of mutations are averaged across all genetic backgrounds at other sites. It has reduced sensitivity to the idiosyncrasy around any particular sequence, but a choice of arbitrary reference state is still required for each site. Its implementation also requires large Hadamard matrices, with the multi-state formalism only recently derived^34^.

Existing methods to address nonspecific epistasis also have limitations. Sometimes the protein’s phenotype can be measured or transformed onto a scale that is expected to be less affected by nonspecific epistasis, such as thermodynamic free energy^16,35,36^. But protein phenotypes can scale nonadditively because of many causes, and the transformation required to remove nonspecific epistasis are seldom known in advance^37^. Even free energy must be measured using techniques that have limited dynamic range and thus entail nonlinearity. Several studies have addressed this issue by inferring a transformation that maximizes the fit of an additive genetic model^9,11,13,17,23,38,39^, but many of these approaches rely on rigid convex or concave transformations that cannot incorporate common forms of nonlinearity, such as the bounding of measurement within lower and upper limits. Some studies employed flexible splines or neural networks^9,23,38^, but these approaches have not been widely adopted because they are cumbersome to implement and interpret.

Here we develop a simple and powerful reference-free framework that is applicable to any number of states, and couple it with an effective model of nonspecific epistasis. We first explain our approach and compare it with existing frameworks. We then systematically reanalyze available combinatorial mutagenesis datasets to assess the complexity of sequence-function relationship. Finally, we explore strategies to infer the genetic architecture when only a small fraction of possible sequences can be experimentally characterized.

## RESULTS

### Reference-free analysis of genetic architecture

We have several goals in dissecting a protein’s genetic architecture. First, we would like to understand the causal rules by which sequence determines function, including the effect of each amino acid at each site, the epistatic interactions between these states, and any global nonlinearities that shape the sequence-function relationship. By decomposing the genetic architecture into its causal components in this way, we can identify the major genetic and biochemical factors that shape a protein’s function. Second, we would like to use these fine-scale causal rules as a foundation for macroscopic descriptions of the genetic architecture, such as the overall importance of effects at each epistatic order or of genetic variation at each site or set of sites. Third, an understanding of the rules of genetic architecture inferred from a sample of variants could allow us to predict the function of any uncharacterized variant.

Our reference-free analysis (RFA) is structured to achieve these goals. RFA treats sequence states, not mutations, as causal factors in a protein’s genetic architecture. This framework allows the genetic causes of phenotypic variation to be defined for the entire ensemble of sequences by adopting Fisher’s statistical framework for dissecting the global effects of genetic states^40^. In RFA, the phenotype of a sequence is the simple sum of the effects of its genetic states (Fig. 1a). The zero-order term, which affects all sequences, is the mean phenotype across sequence space. The additive effect of a state is the difference between the mean phenotype of all sequences containing that state and the global mean. The epistatic effect of a combination of states is the difference between the mean phenotype of all sequences containing the combination and the expected phenotype given the lower-order effects.

**Fig. 1.**
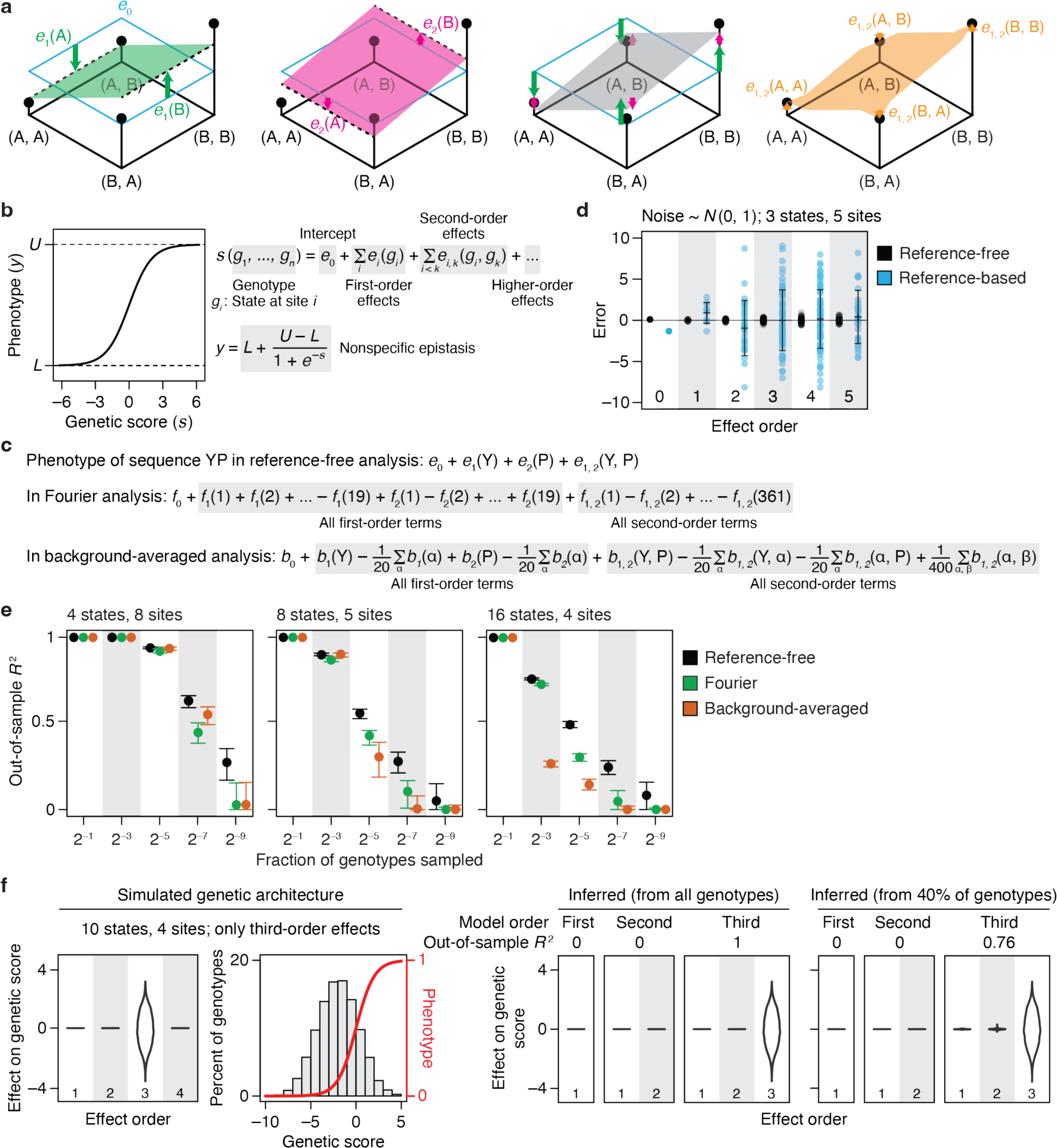
Reference-free analysis (RFA) of genetic architecture. **a**, Illustration of RFA on a 2-state, 2-site genotype space. (*First panel*) The four possible genotypes (dots) are arranged on a plane with the phenotype of each indicated by elevation. The zero-order term (*e*_0_) is the mean phenotype of all genotypes, marked by the horizontal cyan plane. The additive effect of state A or B at site 1 [*e*_1_(A) or *e*_1_(B), green arrows] measures how the mean phenotype of all genotypes containing that state (dashed line) differs from the global mean. The green plane predicts the phenotype based on state at site 1. *(Second panel*) Additive effects at site 2 [*e*_2_(A) and *e*_2_(B)] are defined similarly and represented with pink arrows and plane. *(Third panel)* The complete additive model predicts the phenotype as the sum of the additive effects of all sequence states plus the global mean, represented as the grey plane tilted in both dimensions. (*Fourth panel)* The pairwise interaction between states A and B at sites 1 and 2 [*e*_1, 2_(A, B)] measures how the mean phenotype of all genotypes containing the two states [here just one genotype (A, B)] differs from the additive prediction. **b**, We implement RFA with a sigmoid link function to incorporate nonspecific epistasis. Each variant’s genetic score (*s*) is the net effect of its genetic states. The link function transforms *s* of each variant into its phenotype, *y*. Parameters *L* and *U* of the sigmoid represent the lower and upper bound of measurable phenotype. **c**, Mapping of genetic effects to phenotype in reference-free, Fourier, and background-averaged analyses, shown for an example two-amino acid sequence YP. *e,* reference-free effects; *f_i_*(*k*), the *k*-th Fourier coefficient for site *i*; *b,* background-averaged effects for each possible amino acid state (α) and pair (α, β). **d**, RFA is robust to measurement noise. For a 3-state, 5-site genotype space, reference-free and reference-based effects were computed from phenotype data simulated by sampling measurement noise from the standard normal distribution. Each dot shows the error for an effect at the specified order, and error bars indicate their standard deviation. **e**, RFA is more robust to missing genotypes than are Fourier and background-averaged analyses. Genetic architectures with the indicated number of states and sites were simulated by drawing up to third-order reference-based effects from the standard normal distribution. Third-order models were fit to a varying fraction of genotypes and evaluated by predicting the phenotypes of the remaining genotypes. The mean and 95% interval of out-of-sample *R*^2^ across 200 trials are shown. **f**, RFA does not misinterpret high-order epistasis as clusters of low-order interactions. Phenotypes were simulated with only third-order determinants (distribution shown in the first panel) under a sigmoid link function (second panel). First-, second-, and third-order reference-free effects were inferred from all genotypes or a random subset. The distribution of inferred effects and the variance they explain are shown in the right panels.

This way of dissecting the sequence-function relationship gives RFA several desirable properties. First, it provides an accurate bird’s-eye view of the effects of sequence states and combinations across the entire space of possible variants. By contrast, reference-based analysis (RBA) treats mutations as the determinants of phenotype, which can cause the inferred genetic architecture – including its apparent complexity – to vary wildly depending on the choice of wild-type sequence (see Supplementary Text for illustration). RBA’s description of the genetic architecture also implies that sequences with wild-type residues have no genetic determinants involving those sites. For example, the wild-type sequence has no genetic determinants whatsoever because it contains no mutations. A point mutant is subject to the additive effect of one mutation but is by definition unaffected by epistasis. A double mutant is shaped by one pairwise interaction but no higher-order epistasis, and so on. In reality, these proteins have a genetic architecture just as interesting and potentially complex as that of sequences distant from the wild-type. RBA can therefore be suited to understanding the effects of mutations in the local neighborhood of a particular reference sequence, but it is less useful than RFA for understanding the sequence-function relationship per se.

A second advantage of RFA is that its terms map to phenotype in a straightforward way: a variant’s phenotype is simply the sum of the model terms for the amino acid states and combinations in its sequence, and each term has a simple and intuitive genetic meaning – the average phenotypic effect of the state or combination relative to the global mean. By contrast, Fourier analysis (FA) and background averaging (BA) make it difficult to interpret from the model terms how the function of any variant arises from its sequence. In FA, the phenotype of any sequence is a weighted sum of every single term in the model (Fig. 1c), each of which expresses the effect of a recoded Fourier dimension, not that of an amino acid; when computed for datasets with more than two states per site, these terms have no straightforward genetic or biochemical meaning. BA phenotypes are also a sum over every term in the model, and each term is defined relative to an arbitrary reference state.

Third, RFA offers a maximally efficient description of the global genetic architecture. The RFA model truncated at any epistatic order captures the maximum amount of phenotypic variance that can be captured by any linear model of the same order (see proofs in Supplementary Text; this property is shared with FA and BA but not RBA). Consider the zero-order models, which predict the phenotype of every sequence by a single number. The zero-order term is the mean phenotype of all sequences, which represents the best possible predictor because it minimizes the total squared error. In first-order RFA, the predicted phenotype of a sequence is the sum of the additive effects of its constituent states plus the global mean. This predictor again achieves the minimum total squared error among all possible first-order predictors and therefore explains the maximum possible amount of phenotypic variance. In contrast, the term in a zero-order RBA model – the wild-type phenotype – predicts the wild-type sequence with perfect accuracy but is less accurate across sequence space as a whole. Similarly, the terms in the first-order model – the effect of each point mutation when introduced into the wild-type – perfectly predict single-mutant variants, but across the vast number of other sequences they leave more variation to be explained by epistasis.

Finally, RFA allows direct quantification of the amount of phenotypic variation caused by any set of states or interactions. The phenotypic variance attributable to a reference-free effect is simply the square of its magnitude normalized by the fraction of sequences that involve the term (Supplementary Text). The contribution of a set of model terms, such as the effects of all states at some sites or all interactions at an epistatic order, is simply the sum of the individual contributions. This contrasts with RBA, BA, and FA, in which the terms themselves have no direct relationship to the variance explained.

Although the coefficients of any of the four formalisms can be converted into those of the others using a linear re-mapping, RFA is therefore unique in that its structure directly expresses the rules of genetic architecture in a way that achieves maximal accuracy, efficiency, and interpretability.

Nonspecific epistasis can be incorporated into RFA by using a generalized linear model in which the phenotype of a sequence is a nonlinear transformation of the effects of its genetic states (Fig. 1b). Here we use a simple sigmoid link function defined by two parameters – the minimum and maximum observable phenotype – to account for the diminishing effects of genetic states near the limits of dynamic range; these parameters and the specific terms of the RFA model can be estimated jointly by least-squares regression. When this function is applied, each term in the RFA model reflects the average effect of the state or combination on a variant’s genetic score – the sum of all the states and combinations in its sequence – and the phenotype is the nonlinear function of the score.

### Robustness to measurement noise, missing genotypes, and model misspecification

If precise phenotypic measurements were available for every sequence, the causal factors as defined by any of the formalisms could be computed exactly. But real experiments are subject to noise, and some variants are usually missing. Another challenge is that when there are many variable sites, the complete model contains too many parameters to estimate with confidence, so truncated low-order models are typically used for inference. These models are misspecified if there are higher-order interactions in the data, potentially leading to biased inference^27^. RFA is designed to be robust to these challenges.

RFA is robust to noise because each effect term is computed using average measurements over many variants. In contrast, each effect in an RBA model is computed as a chain of sums and subtractions of individual variants, so the measurement errors propagate and snowball with epistatic order. To compare the performance of RFA and RBA in the face of measurement noise, we simulated phenotypic data using a known genetic architecture with normally distributed noise. We computed the effect terms from these data and compared them to the true values (Fig. 1d). RFA effects are estimated precisely, with an average error much less than that for individual phenotypic measurements. By contrast, the errors in the estimates of RBA effects are much greater than that for individual genotypes, and they increase dramatically with epistatic order.

When sampling of variants is incomplete, least-squares regression can be used to accurately estimate RFA effects. RFA model terms are defined using mean phenotypes over sets of variants, so the least-squares estimates are unbiased as long as sequences are sampled at random, and they converge to the true RFA effects as more sequences are sampled (Supplementary Text). RBA terms, by contrast, are defined as differences between particular individual genotypes, which have no expected relationship to the least-squares estimates. Using simulated genetic architectures, we found that the least-squares regression estimate of RBA effects are inaccurate and explain more phenotypic variance than the true RBA effects do, unless the model contains all the nonzero terms in the genetic architecture.^27^ The mismatch between the true RBA architecture and that estimated by least-squares regression becomes very severe in the presence of noise and when genotypes are missing (Supplementary Text).

FA and BA models can be estimated by regression because their terms are defined as averages, but they are parameterized differently from RFA, so the accuracy of regression-estimated terms could differ. We simulated a genetic architecture, removed a variable fraction of sequences, fit the models to the remaining sequences, and used the inferred models to predict the phenotypes of the excluded sequences (Fig. 1e). When there are only four possible states per site, all models have high predictive accuracy, declining only after the fraction of sampled sequences drops below 1%, at which point RFA is slightly more accurate. When there are 16 states, however, RFA is much more robust than BA, the accuracy of which degrades rapidly as the sample size shrinks; it is also more robust than FA but to a less extent. RFA is more robust to missing genotypes because the phenotype of an unsampled variant is predicted as the sum of only the terms for its genetic states. FA and BA predict the phenotype as a weighted sum of all model terms (Fig. 1c), so the error associated with each term propagates to every genotype. This difference is exacerbated as more states are considered, because the total number of terms increases exponentially with the number of states.

Finally, we explored the possibility that RFA might oversimplify the genetic architecture by misinterpreting high-order interactions as clusters of lower-order effects, particularly when truncated models and a link function to account for nonspecific epistasis are used. RFA is structured so that each order of terms produces a distinct pattern of phenotypic variation, and the pattern produced by terms at one order cannot be explained by terms of a different order. High-order variation appears as noise around the mean at lower orders, so a truncated RFA model in principle cannot explain any phenotypic variation caused by unmodeled higher-order interactions (Supplementary Text). To test whether this expectation holds in practice, we simulated phenotypes under a genetic architecture that contains only third-order effects plus nonspecific epistasis and then fitted RFA models (with the sigmoid link function) truncated at various orders (Fig. 1f). As expected, the first- and second-order truncated models correctly explain zero phenotypic variance and detect no first- or second-order effects. When the third-order model is used, all variance is correctly attributed to third-order interactions. Similar results obtain when variants are only partially sampled. Neither model truncation nor the use of a nonlinear link function therefore causes artifactually simple inferences of complex genetic architecture under these conditions.

These analyses establish that the effect terms and the variance partition of RFA can be estimated accurately from noisy and partial data, whereas those under the RBA formalism cannot be. FA and BA share some of RFA’s advantages but are less accurate when inferring architectures with many possible states from incomplete data.

### Simplicity of protein sequence-function relationships

To understand the genetic architecture of real proteins, we performed RFA on 20 combinatorial mutagenesis datasets available for antibodies, enzymes, fluorescent proteins, transcription factors, viral surface proteins, and toxin-antitoxin pairs (Table 1). We considered only datasets with precise measurement (*r*^2^ > 0.9 among replicates) and sampling of at least 40% of possible variants. We focused primarily on large libraries but included three small ones in which high-order epistasis has been reported. The datasets range in size from 32 to 160,000 possible genotypes, with the number of variable sites ranging from 3 to 16 and the number of sampled states per site from 2 to20. To assess the complexity of genetic architecture, we fitted to each dataset a series of truncated reference-free models of increasing order, using the sigmoid link function to incorporate nonspecific epistasis and L1 regularization to reduce overfitting; we then used cross-validation to estimate the fraction of phenotypic variance explained at each model order as the out-of-sample *R*^2^.

**Table 1.** Combinatorial mutagenesis datasets analyzed in this study.

| Protein | Genotype space | Phenotype |
| --- | --- | --- |
| Methyl-parathion hydrolase<sup>43</sup> | 2<sup>5</sup> (32) | Catalytic activity |
| β-lactamase<sup>45</sup> | 2<sup>5</sup> (32) | Antibiotics resistance (MIC) |
| Dihydrofolate reductase<sup>3</sup> | 3 × 2<sup>4</sup> (48) | Antibiotics resistance (IC<sub>75</sub>) |
| Influenza A H3N2 hemagglutinin<sup>39</sup> | 2<sup>2</sup> × 3<sup>2</sup> × 4<sup>2</sup> (576) | Viral replication fitness |
| Antibody CR6261<sup>17</sup> | 2<sup>11</sup> (2,048) | Affinity for influenza hemagglutinin strain H1 or H9 |
| Bacterial antitoxin ParD3<sup>46</sup> | 20<sup>3</sup> (8,000) | Fitness conferred by binding to toxin ParE3 or ParE2 |
| *Aequorea victoria* GFP (avGFP)<sup>13</sup> | 2<sup>13</sup> (8,192) | Fluorescence |
| Bacterial antitoxin ParD3<sup>47</sup> | 13 × 12 × 10 × 6 (9,360) | Fitness conferred by binding to toxin ParE3 or ParE2 |
| SARS-CoV-2 spike protein<sup>7</sup> | 2<sup>15</sup> (32,768) | Affinity for human ACE2 |
| Antibody CH65<sup>18</sup> | 2<sup>16</sup> (65,536) | Affinity for influenza hemagglutinin strain MA90, MA90-G189E, or SI06 |
| Antibody CR9114<sup>17</sup> | 2<sup>16</sup> (65,536) | Affinity for influenza hemagglutinin strain B, H1, or H3 |
| Transcription factor ParB<sup>44</sup> | 20<sup>4</sup> (160,000) | Fitness conferred by transcription |
| Protein G B1 domain (GB1)<sup>10</sup> | 20<sup>4</sup> (160,000) | Binding enrichment for IgG-Fc |

We found that genetic architecture is simple, with most phenotypic variance explained by main effects of amino acids and virtually all of the remainder by pairwise interactions. The first-order model achieves a median *R*^2^ of 0.91 across the 20 datasets – with a maximum of 0.97 and *R*^2^ > 0.75 in all but four cases (Fig. 2a). There is no relationship between the fraction of variance explained at the first order and the number of sites or states assayed (Extended Data Fig. 1). When pairwise interactions are included, virtually all genetic variance is explained, with a median out-of-sample *R*^2^ of 0.96 and a minimum of 0.92 across the datasets.

**Fig. 2.**
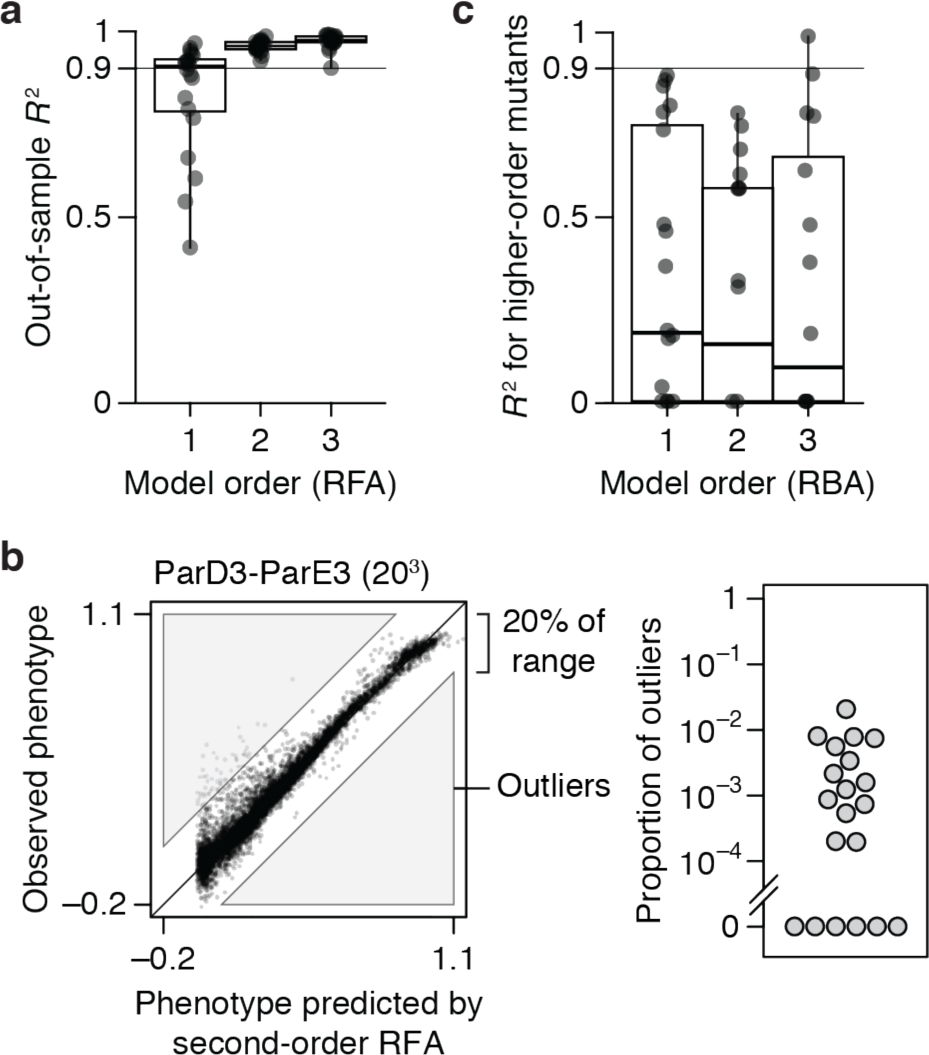
Simplicity of protein sequence-function relationships. **a**, RFA of 20 combinatorial mutagenesis datasets (Table 1). First-, second-, and third-order models with the sigmoid link function were evaluated by cross-validation – by inferring the model from a subset of data and predicting the rest of data. Each dot shows the mean out-of-sample *R*^2^ for one dataset; boxplots show the median, interquartile range, and total range across the datasets. Extended Data Fig. 1 shows the *R*^2^ for individual datasets. **b**, Variants possibly affected by strong high-order epistasis were identified as outliers in the second-order model (residual greater than 20% of the phenotype range). (*Left*) Outliers in the ParD3-ParE3 (20^3^) dataset. Each point is a variant, plotted by its observed and predicted phenotype. (*Right*) Proportion of outliers in each dataset. **c**, Reference-based analysis of the 20 datasets. Each model was evaluated by predicting the phenotypes of higher-order mutants. Nonspecific epistasis was accounted for as in **a**, and the genotyped designated as wild-type in the original publication was used as reference. Negative *R*^2^ values are plotted as zero.

Incorporating third-order terms offers only marginal or no improvement in fit (median change in out-of-sample *R*^2^ of 0.02, maximum of 0.04). The small fraction of phenotypic variance unexplained by the third-order model represents some combination of fourth- and higher-order epistasis, measurement noise, and limitations in the sigmoid link function to accurately capture nonspecific epistasis. These results are not attributable to the use of regularization (Extended Data Fig. 2).

Although high-order epistasis is negligible across sequence space as a whole, there could still be a subset of genotypes shaped by strong high-order epistasis. To address this possibility, we analyzed the residuals of the second-order model, which are the sum of all higher-order interactions and measurement noise. Genotypes with a residual greater than 20% of the phenotype range were considered candidates for strong higher-order epistasis, although erratic measurement noise cannot be excluded. The proportion of such genotypes is zero in six datasets and between 0.02 to 2% in the others (Fig. 2b). Only a tiny fraction of genotypes is therefore potentially affected by strong high-order epistasis.

These analyses show that the genetic architecture of proteins is simple: knowing just the additive effects and pairwise interactions, coupled with a simple model of nonspecific epistasis, is sufficient to accurately account for and predict the phenotype across the entire ensemble of sequences. Higher-order interactions are not completely absent, but they are weak and limited to a very small fraction of genotypes.

Finally, we also examined the 20 datasets by RBA. We fitted first-, second-, and third-order reference-based models (along with the sigmoid link function) to the wild-type sequence and all its single, double, and triple mutants. The fitted models were then evaluated by predicting the phenotypes of higher-order mutants. The median *R*^2^ across datasets is less than 0.2 at any order (Fig. 2c), leaving the vast majority of phenotypic variance to be explained by fourth- or higher-order epistasis. RBA therefore leads to unnecessarily complex estimates of protein genetic architecture.

### The primary cause of nonspecific epistasis is phenotype bounding

To understand the impact of incorporating nonspecific epistasis, we compared RFA of the empirical datasets with and without the sigmoid link function. We found that incorporating nonspecific epistasis dramatically improves phenotype prediction and reduces the variance attributed to epistasis (Fig. 3a, b). Using the sigmoid link raises the median out-of-sample *R*^2^ of first-order models from 0.59 to 0.92, reducing the variance that must be explained by epistasis by a factor of 5. For second-order models, it improves the median *R*^2^ from 0.87 to 0.96, reducing the variance attributed to higher-order epistasis by a factor of three. For third-order models, incorporating nonspecific epistasis increases the median *R*^2^ from 0.95 to 0.98.

**Fig. 3.**
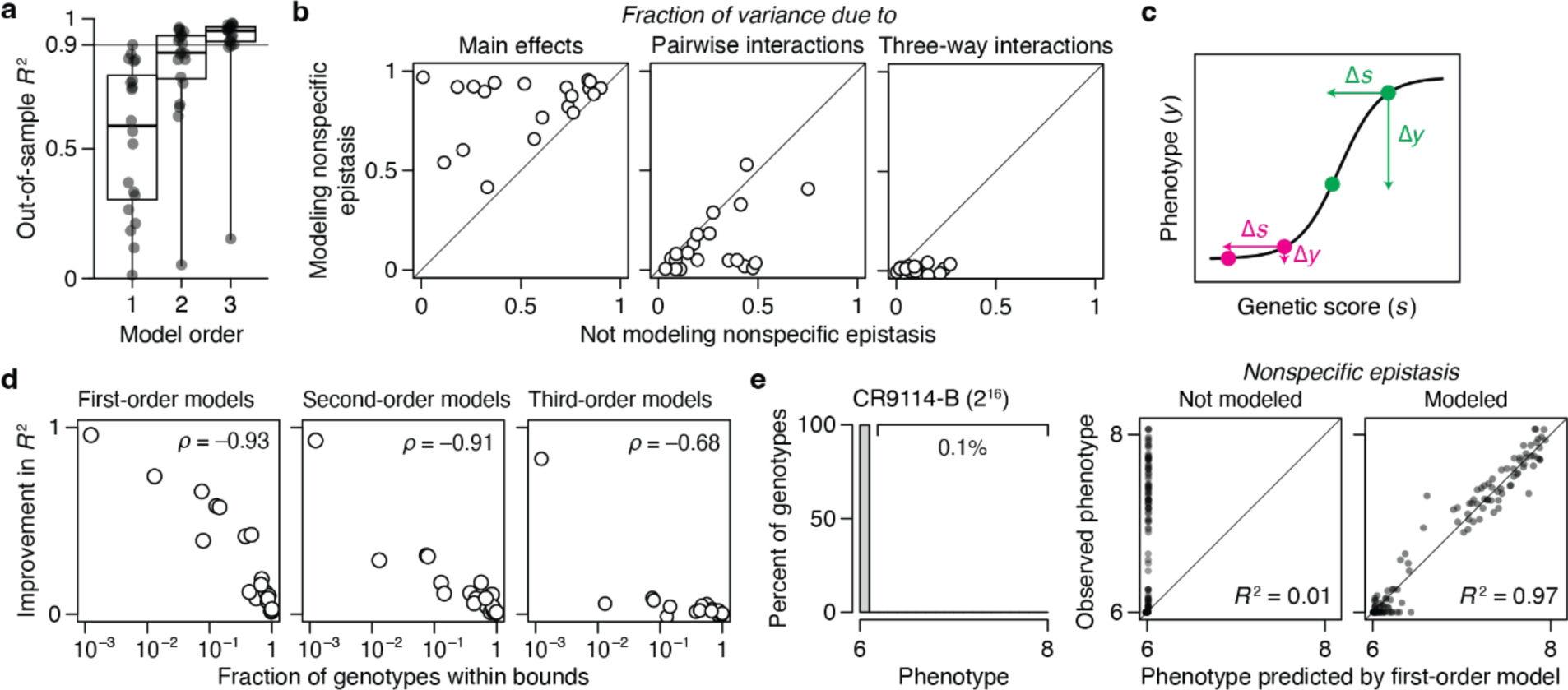
The primary cause of nonspecific epistasis is phenotype bounding. a, RFA of the 20 datasets without incorporating nonspecific epistasis, shown as in Fig. 2a. **b**, Incorporating nonspecific epistasis reduces the amount of phenotypic variance attributed to pairwise and higher-order interactions. Each dot shows the variance component for one dataset computed with or without incorporating nonspecific epistasis. **c**, Nonspecific epistasis causes the phenotypic effect of a mutation (Δ*y*) to vary among genetic backgrounds (magenta versus green) even when the effect on genetic score (Δ*s*) is constant. Phenotype bounding is a particularly strong form of nonspecific epistasis that causes mutations to appear neutral on backgrounds near the bounds but not on others. **d**, The extent to which the sigmoid link function improves the model fit (comparing out-of-sample *R*^2^ in Fig. 3a versus 2a) is proportional to the fraction of genotypes at the phenotype bounds. **e**, In a dataset where only 0.1% of genotypes are within the bounds, incorporating nonspecific epistasis raises the fraction of phenotypic variance explained by additive effects from 0.01 to 0.97.

The dramatic improvement in fit conferred by the simple sigmoid function suggests that phenotype bounds – biological or technical limits on the dynamic range over which genetic states have measurable effects on function – are the primary cause of nonspecific epistasis (Fig. 3c). Corroborating this conclusion, the degree to which the link function improves the *R*^2^ is tightly correlated with the fraction of genotypes at the phenotype bounds (Fig. 3d). In the CR9114-B dataset, for example, 99.9% of genotypes are at the lower bound, and incorporating nonspecific epistasis improves the first-order variance explained from 1% to 97% (Fig. 3e). Although the causes of nonspecific epistasis are likely to be complex and vary among datasets, the simple sigmoid therefore captures its most salient features.

### Sparsity of protein sequence-function relationships

We next asked whether protein function is determined by many genetic states and interactions of small effect or by a few determinants of large effect. For each dataset, we estimated the minimal number of reference-free terms required to predict the function with 90% accuracy (*T*_90_): we ranked the terms in the fitted third-order model by their contribution to variance, constructed increasingly complex models by sequentially including each term, and estimated the accuracy of each model by cross-validation (Fig. 4a).

**Fig. 4.**
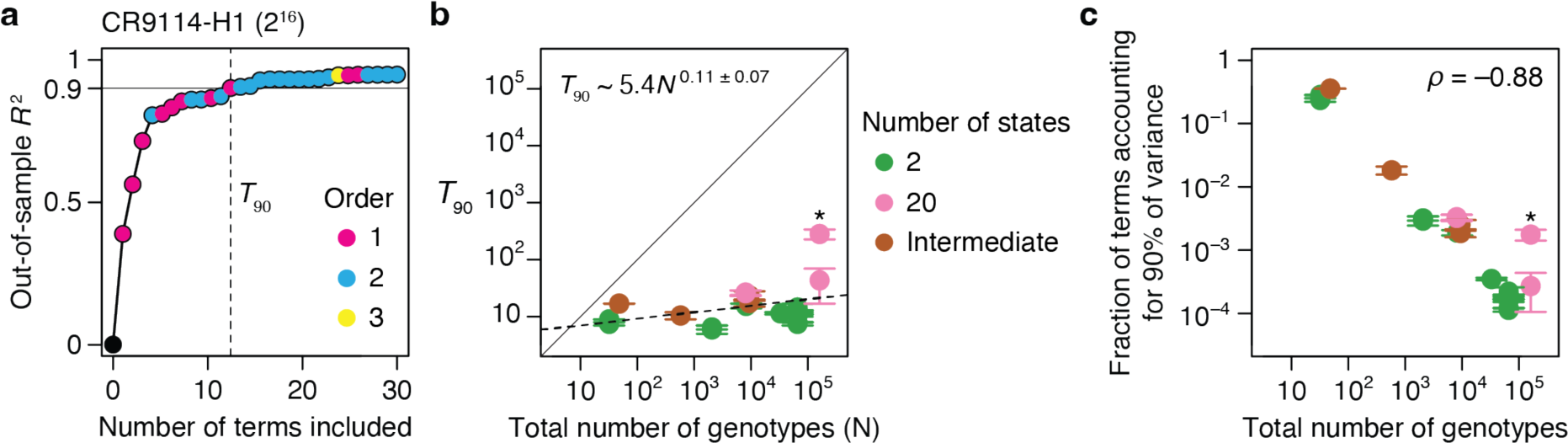
Sparsity of protein sequence-function relationships. **a**, Measuring the sparsity of genetic architecture illustrated on the CR9114-H1 dataset. Up to third-order reference-free effects were estimated and ranked by the fraction of variance they explain. Models of increasing complexity were constructed by sequentially including each effect term, and each model was evaluated by cross-validation. Each dot represents a model, colored by the order of the last term added. Vertical line marks *T*_90_, the minimal number of terms required for an out-of-sample *R*^2^ of 0.9. **b**, *T*_90_ as a function of the total number of genotypes. Dotted line, best-fit power function. Asterisk, GB1 dataset. Each *T*_90_ was estimated in two ways: as the number of terms required to reach *R*^2^ of 0.9 (upper error bar) – an overestimate because measurement noise prevents any model from attaining an out-of-sample *R*^2^ of 1 – and as the number of terms required for an *R*^2^ equal to 90% of that of the full third-order model (lower error bar). Circles show the average of the two estimates. **c**, Fraction of all terms required to explain 90% of phenotypic variance shown against the total number of genotypes. Asterisk, GB1 dataset.

The genetic architecture of proteins is very sparse (Fig. 4b). Out of up to 160,000 possible terms, *T*_90_ ranges from just 6 to 44 across all datasets except for GB1, in which the mutated sites were specifically chosen to be enriched for epistatic interactions^10^. As the total number of possible genotypes (*N*) increases, *T*_90_ increases very slowly, so that the fraction of all terms required for an *R*^2^ of 0.9 declines almost linearly (Fig. 4c). These relationships hold irrespective of the number of states per variable site.

Our findings suggest that even a very large genetic architecture should be describable with a compact set of terms. For example, the relationship between *T*_90_ and *N* predicts that a very large genetic architecture – two states at 100 variable sites (∼10^30^ possible genotypes and model terms) – could be described with 90% accuracy by a model with just ∼10,000 key terms.

### Inferring genetic architecture by sparse sampling

Although a protein’s genetic architecture is defined by relatively few causal factors, identifying them could be challenging. Comprehensive experimental characterization is impractical for sequence spaces much larger than those we have analyzed, so a critical question is whether the important terms can be inferred from a small sample of genotypes by sparse learning methods^13^. To address this possibility, we sampled a variable number of genotypes from the datasets, fitted reference-free models using regression with L1 regularization, predicted phenotypes of the unsampled genotypes, and determined *N*_90_, the minimum sample size required for *R*^2^ of 0.9 (Fig. 5a).

**Fig. 5.**
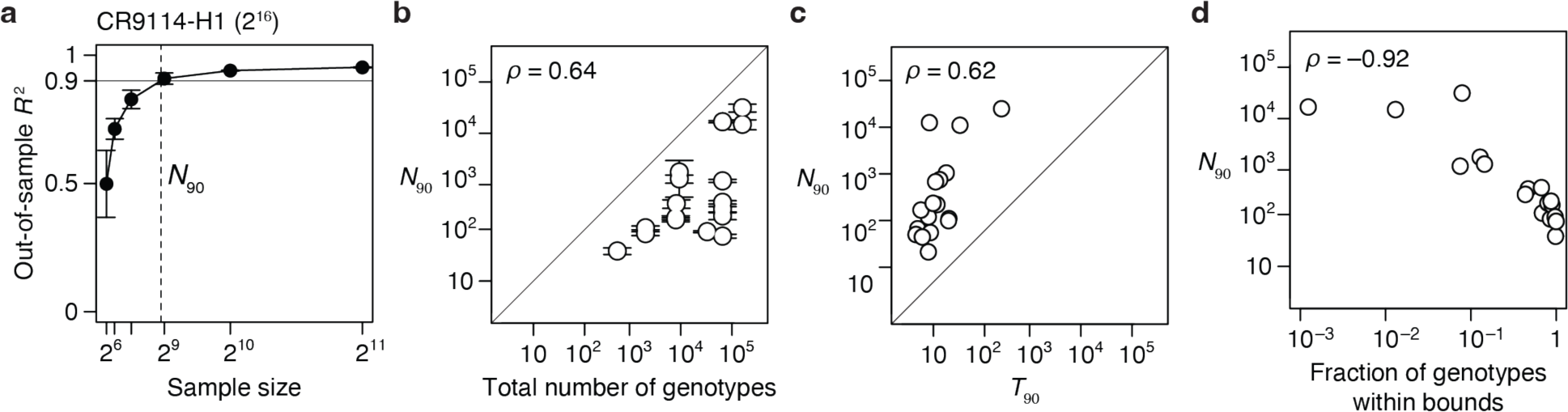
Inferring genetic architecture by random sampling. **a**, Inference illustrated on the CR9114-H1 dataset. Up to third-order reference-free effects were inferred from a varying number of randomly sampled genotypes, and were evaluated by predicting the phenotypes of all unsampled genotypes. For each sample size, the mean and standard deviation of out-of-sample *R*^2^ across 10 trials are shown. Dashed line marks *N*_90_, the minimal sample size required for a mean out-of-sample *R*^2^ of 0.9. **b**, *N*_90_ as a function of the total number of genotypes. Error bars were computed as in Fig. 4a. The three datasets with 48 or fewer genotypes are not shown. **c**, *N*_90_ as a function of *T*_90_, the minimal number of terms required to explain 90% of phenotypic variance (Fig. 4). **d**, *N*_90_ as a function of the fraction of genotypes within phenotype bounds.

We found that genetic architecture of proteins cannot be efficiently inferred from sparse random samples (Fig. 5b). Excluding the three small datasets, *N*_90_ ranges from 0.2 to 25% of the total number of genotypes, with a median of 5%. Even the lowest end of this range does not bode well for inferring the architecture of large sequence spaces with many states at many variable sites.

We evaluated several factors that might determine the necessary sample size. First, large sequence spaces require larger samples: *N*_90_ increases with the total number of genotypes, although there is a considerable scatter in this relationship (Fig. 5b). The complexity of the genetic architecture is not a major factor: *N*_90_ depends only weakly on *T*_90_ (Fig. 5c). A critical factor is the fraction of genotypes within the dynamic range of measurement: *N*_90_ increases sharply with the degree of phenotype bounding (Fig. 5d). An extreme case is the CR9114-B dataset (65,536 genotypes), where just 10 additive effects account for 90% of phenotypic variance but >16,000 genotypes are needed to identify them. This is because 99.9% of genotypes are at the lower bound, providing little quantitative information on genetic effects. By contrast, the CH65-MA90 dataset consists of the same number of genotypes, but the genetic architecture can be inferred from just 99 random genotypes because there is virtually no phenotype bounding.

We conclude that despite the global simplicity of proteins’ genetic architecture, the important causal factors cannot be efficiently identified by sparse random sampling. A critical step is therefore to develop a non-random sampling strategy that can efficiently identify the key additive effects and pairwise interactions defining a genetic architecture.

### Understanding genetic architecture

A benefit of coupling RFA with the sigmoid link function is that the genetic effects are expressed in a unit that is intelligible through a simple biophysical analogy, and they become comparable across datasets, even when different phenotypes are measured. The sigmoid model describes the phenotype of a variant as an equilibrium between two thermodynamic states: the functional state, whose phenotype is *U*, and the nonfunctional state, whose phenotype is *L* (Fig. 6a). A variant’s phenotype, lying between *U* and *L*, reflects the relative occupancy of the functional to nonfunctional state, which is determined by its genetic score (*s*) as *e^s^*. The genetic score takes the role of the Gibbs free energy difference between the two states (Δ*G*) expressed in the unit of –*kT*. If a variant’s genetic score is 0, the two states are equally populated and its phenotype is midway between *U* and *L*. A sequence state or combination that increases the genetic score by 2.3 always causes a ten-fold increase in the relative occupancy of the functional state, corresponding to a ΔΔ*G* of –1.4 kcal/mol at 37°C. This relationship holds across proteins, functions, and experimental systems.

**Fig. 6.**
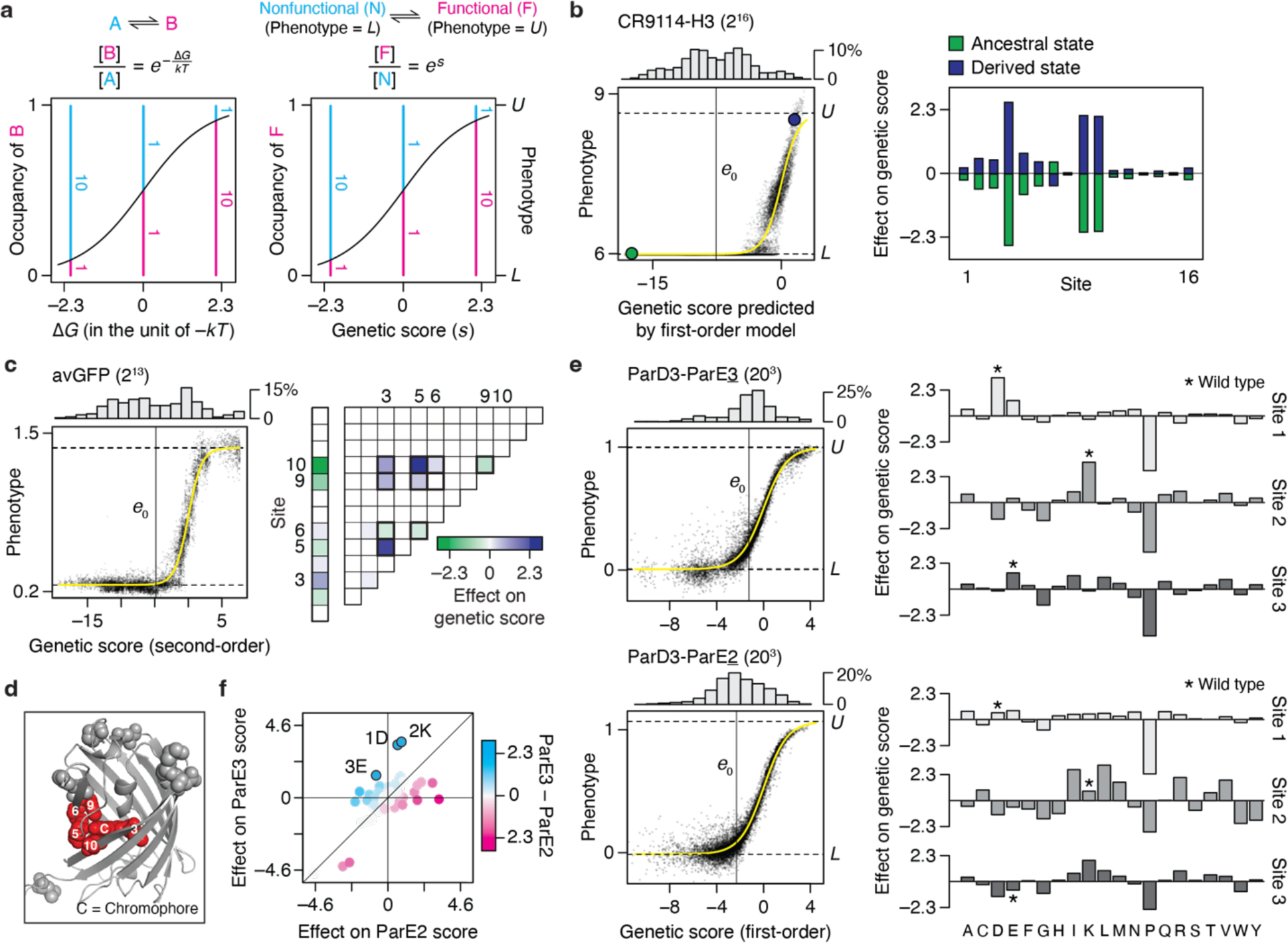
Understanding genetic architecture. **a**, Interpreting genetic score (*s*) as free energy difference (Δ*G*). (*Left*) Relative occupancy of two thermodynamic states as a function of their Δ*G*. *k*, Boltzmann constant; *T*, absolute temperature. (*Right*) The sigmoid link function can be interpreted as describing an equilibrium between two states—the “functional” state, of phenotype of *U*, and the “nonfunctional” state, of phenotype of *L*. Their relative occupancy (pink versus blue lines) equals *e^s^*, allowing *s* to be interpreted as Δ*G* in the unit of –*kT*. **b**, CR9114-H3 dataset, which measures the affinity of all combinations of ancestral and derived amino acids at 16 sites in an antibody towards a hemagglutinin. (*Left*) First-order RFA. Each dot is a genotype, plotted by its measured phenotype and estimated genetic score. Histogram, distribution of genetic score; yellow curve, inferred sigmoid link; horizontal lines, inferred phenotype bounds; vertical line, mean genetic score; green and purple dots, ancestral and derived genotypes. (*Right*) Additive effects of amino acids at each site. **c**, avGFP dataset, which measures the fluorescence of all combinations of pairs of amino acids at 13 sites. (*Left*) Second-order RFA. (*Right*) Additive effects and pairwise interactions, which account for 57 and 38% of phenotypic variance, respectively. Values are shown for one of the two of amino acids at each site. The ten pairwise interactions possible among sites 3, 5, 6, 9, and 10 are outlined. **d**, Crystal structure of avGFP (PDB ID: 3e5w). The 13 mutated sites are shown in spheres, and the chromophore and its five surrounding sites are colored in red. **e**, ParD3-ParE3 and ParD3-ParE2 (20^3^) datasets, which measure how all possible variants of ParD3 at three sites bind to ParE3, the cognate substrate, or ParE2, a noncognate substrate. (*Left*) First-order RFA. (*Right*) Additive effects at each site. Asterisk, wild-type amino acid. **f**, Comparing the effect of each amino acid on ParE3 versus ParE2 binding. Wild-type amino acids are marked.

We applied this framework to understand the genetic architecture of several example proteins. The CR9114-H3 dataset (Fig. 6b) consists of affinity measurements for binding of 2^16^ antibody variants (all possible combinations of ancestral and derived amino acids at 16 sites that evolved during affinity maturation) to an influenza hemagglutinin. The vast majority of variants are at the lower bound of detectable binding, so the average genetic score is –7.8, corresponding to just 0.04% occupancy of the bound state. The best variant has a score of just 2.6, corresponding to 93% occupancy. There is virtually no specific epistasis in this genetic architecture (Extended Data Fig. 1). Additive effects at three key sites mostly determine the phenotype: each favorable state increases the genetic score by 2.1 to 2.6, together increasing the relative occupancy by almost three orders of magnitude but still yielding absolute occupancy of the bound state at just 36%. The state at five other sites make modest contributions, each changing the genetic score by around 0.5 and shifting the relative occupancy by ∼25%; the remaining eight have even smaller effects. A variant must therefore have all three large-effect favorable states to achieve measurable binding, and the particular combination of states at the other sites modulates the affinity. This architecture reflects and explains why most variants are at the lower bound of measurement, why the global average genetic score and occupancy of the bound state are so low, and why the maximum observed affinity and fraction bound are very modest.

Specific pairwise interactions are important in the avGFP dataset (Fig. 6c), accounting for 38% of variance in fluorescence measurements. There are many functional variants in this library, including a large number at the measurement maximum, which gives the average variant a genetic score ∼–1 and occupancy of the fluorescent state ∼20%. First- and second-order effects involving just five of 13 variable sites account for 86% of variance. These sites, which tightly surround the chromophore in the crystal structure (Fig. 6d), engage in a dense epistatic network in which nine of the ten possible pairwise interactions are nonzero. Although only four of these interactions alter the genetic score by >1, their total impact is substantial, conferring an increase in genetic score by 7.8 and relative occupancy by 2,400-fold in the most favorable genotype. Not all of these are necessary to achieve high fluorescence, however: because only pairwise interactions are involved and the global average has measurable fluorescence, one or more states can be removed while leaving the other favorable interactions intact.

RFA terms can also be used to understand the determinants of functional specificity in multistate sequence space and when multiple functions are measured. The ParD3 library (all combinations of 20 states at 3 sites in the binding interface) was assayed separately for binding its cognate ligand ParE3 and the noncognate ligand ParE2; effects on specificity can be quantified as the difference between a state’s genetic score for the two ligands. The average variant displays a weak but measurable binding to both ligands, with a preference for ParE3 over ParE2 by a genetic score of ∼1 (difference in relative occupancy of 2.5-fold). For both ligands, additive effects account for the vast majority of variance in binding (Fig. 6e). Only eight amino acid states can change the genetic score in favor of one ligand over the other by >1.6, each equivalent to a >5-fold difference in occupancy (Fig. 6f). The three strongest of these each favor ParE3 by scores of 2.2 to 2.8 (∼10-fold preference in occupancy) – two by increasing affinity for both ligands but more strongly enhancing ParE3 binding, and the third via opposite effects on the two ligands. The wild-type protein in this case possesses these three specificity-optimal states.

## DISCUSSION

Our finding that additive effects and pairwise interactions account for virtually all genetic variation within proteins contrasts with several reports of extensive high-order epistasis^1–7^. Use of reference-based analysis and incomplete accounting of nonspecific epistasis have led prior studies to invoke more high-order epistasis than is necessary to explain the data.

We expect our finding to be general across proteins and biochemical phenotypes. The datasets we analyzed comprise proteins with diverse structures and functions. It is unlikely that the particular sites varied in the datasets biased the architectures towards simplicity: in most cases, the sites were chosen because of prior structural evidence that they are functionally important or they vary between functional homologs. The sites are dispersed across the structure in some datasets but clustered in others. A limitation is that each dataset assessed a single phenotype, so the genetic architecture of functional specificity could be more complex; however, a recent study using a similar approach as ours found that high-order interactions within a transcription factor are unimportant for determining its DNA binding specificity^41^.

The lack of high-order epistasis within proteins may seem surprising from a structural perspective, because proteins often contain clusters of three or more residues that contact each other directly. Our results indicate that the effects of such clusters can largely be explained as the sum of the pairwise interactions. But any pairwise coupling depends on the fold of the protein, which in turn depends on states at other sites. A mutation that changes the conformation should alter pairwise couplings and induce high-order epistasis. In the datasets we examined, such conformational epistasis seems rare or inconsequential. A possible explanation is that these datasets held constant the majority of sites in the protein and therefore presumably maintained the overall conformation (or they unfolded the protein). High-order interactions that specify a protein’s fold might be revealed in a library large enough to contain variants with distinct conformations.

The effectiveness of the sigmoid link to capture nonspecific epistasis may seem surprising, because nonlinearities in sequence-function relationships can arise from complex biological and technical causes that vary among proteins, phenotypes, and assays. Our results indicate that bounds on the range over which a phenotype can be produced and measured are the primary cause of nonspecific epistasis. Irrespective of the underlying causes, incorporating this nonlinearity using a simple sigmoid allows a low-order RFA model of specific epistasis to capture virtually all phenotypic variation in the datasets. This joint model provides a parsimonious and efficient description of a protein’s genetic architecture.

Our finding that RFA outperforms RBA in providing a compact and accurate description of the global sequence-function relationship does not mean that RBA is never useful. RBA may be appropriate if the object of interest is interactions among a few mutations in the immediate sequence neighborhood of a designated wild-type or ancestral protein. Even in this case, RBA should be used with caution because of its tendency to infer interactions from measurement noise and local idiosyncrasies.

For scientists who would like to understand how proteins work, our findings are reassuring, but they also clarify a challenge. Proteins’ genetic architecture is intelligible: a small fraction of low-order model terms explains most functional variation. It is therefore not necessary to exhaustively characterize complete combinatorial libraries, which would quickly become intractable as the number of sites or states increases. Sparsely sampling at random from such libraries cannot efficiently identify the important genetic determinants, because the sequence space is vast and most random sequences are nonfunctional. Analyzing the effects of low-order combinations of mutations on a single functional protein would not work either, because this approach would be subject to the same kind of errors and idiosyncrasies that plague RBA. An effective strategy may be to perform single- and double-mutant scans using as starting points a diverse set of functional proteins, such as distantly related homologs^42^, while also improving the dynamic range of measurement. The potential of such a practical strategy to efficiently learn the rules of sequence-function relationships has been largely overlooked, presumably because the genetic architecture of proteins was thought to be much more complex than it is.

## METHODS

### Reference-free analysis (RFA)

Here we define RFA and summarize its key properties. Proofs for the properties and detailed comparisons with other formalisms are in Supplementary Text. Scripts and tutorials for performing RFA are on GitHub (github.com/whatdoidohaha/RFA).

Consider a genotype space defined by *q* states across *n* sites. Let ***g*** denote a genotype, *y*(***g***) its phenotype, and *G* the set of all *q^n^* possible genotypes. RFA decomposes the phenotype into the contribution of individual states and their interactions relative to the global mean phenotype, which is denoted

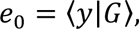

where the bracket notation indicates averaging of *y* over *G*. The additive effect of state *s* in site *i* is the difference between the mean phenotype of the subset of genotypes sharing that state (denoted 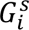) and the global mean:

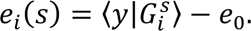

The pairwise interaction between states *s*1 and *s*2 in sites *i*1 and *i*2 is the difference between the mean phenotype of the subset of genotypes sharing that state-pair (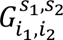) and the global mean after accounting for the additive effects:

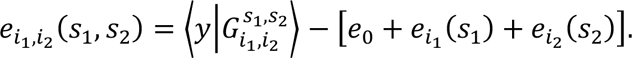

Likewise, a higher-order effect is the difference between the mean phenotype of a subset of genotypes sharing a set of states and the global mean after accounting for the relevant lower-order effects.

RFA predicts the phenotype by summing the effects of all states in the genotype. For a genotype with state *gi* in site *i*, the predicted phenotype under RFA of order *k* is

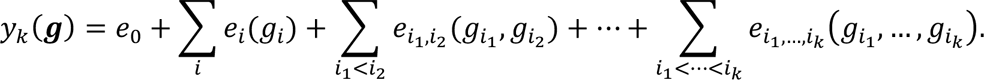

The overall accuracy of this prediction can be quantified by the sum of squared errors

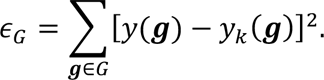

Among all linear models of the same order, including reference-based models under any choice of wild-type genotype, RFA minimizes ε*G* for any *k* for any set of sequence-function associations. For example, when *k* is zero (all phenotypes predicted by a single number), ε*G* is minimized by the global mean phenotype, which is the RFA zero-order term. By minimizing ε*G*, RFA explains the maximum fraction of phenotypic variance that can be explained by any linear model of the same order. Fourier and background-averaged analyses share this property.

RFA facilitates the analysis of genetic architecture by partitioning the phenotypic variance into components attributable to each state and interaction:

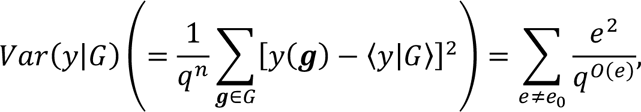

where *e* denotes any nonzero-order effect and *O*(*e*) its order. Note that an effect of order *k* is involved in the phenotype of one in *q^k^* genotypes. The amount of phenotypic variance attributable to an effect is therefore the square of its magnitude normalized by the fraction of genotypes involving that effect.

### Inferring reference-free effects from noisy and incomplete data

When individual phenotypes are subject to measurement noise of variance ω, a reference-free effect of order *k* computed from them has a variance

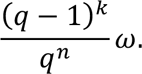

This is always smaller than ω and typically miniscule for low-order effects. The extensive averaging of phenotypic measurements in the computation of reference-free effects confers robustness to measurement noise.

When some genotypes are missing from data, reference-free effects can be inferred by regression. To infer effects of order up to *k*, we model

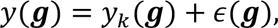

where the residual ε(***g***) is the sum of all higher-order effects and measurement noise. Let *G*\* be the set of sampled genotypes. The regression estimates are obtained by minimizing the sum of squared errors across *G*\*,

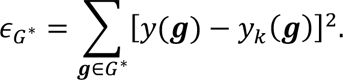

Because reference-free effects minimize the sum of squared errors across genotype space, the regression estimates converge to the true effects as more genotypes are sampled. The estimates are unbiased as long as genotypes are randomly sampled; the unmodeled higher-order effects appear as noise to any lower-order model and therefore do not bias the regression.

### Nonspecific epistasis

We account for nonspecific epistasis by assuming that the effects of sequence states are transformed by a nonlinear link function into the observed phenotype. We modeled the link function as a simple sigmoid, which is defined by two parameters corresponding to the lower (*L*) and upper (*U*) bound of phenotype:

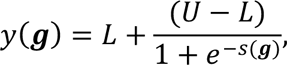

where *s*(***g***) is the sum of the reference-free effects of all states in ***g*** – its genetic score. In principle any link function could be used for this purpose.

### Implementation

We inferred the link function and reference-free effects jointly by regression. The joint inference^23^ is desirable over a widely used two-step approach, which infers the link function first and applies its inverse transformation on the observed phenotype to compute the effects of sequence states^11^. The two-step approach infers the link function by fitting an additive model under the assumption of no nonspecific epistasis and by identifying any systematic nonlinearity between the observed and predicted phenotype. Because the additive model is fit under the incorrect assumption that nonspecific epistasis is absent, this approach cannot uncover the true link function. Furthermore, the inverse transformation can dramatically amplify measurement noise for genotypes near the phenotype bounds.

The joint regression was performed with L1 regularization to reduce overfitting. The optimal regularization strength was determined by maximizing the out-of-sample *R*^2^ in cross-validation. Except for four datasets, cross-validation was performed by randomly partitioning the genotypes into training and test sets. For the three datasets with 48 or fewer genotypes and the CR9114-B dataset where only 81 genotypes are above the lower phenotype bound, cross-validation was performed by leaving out each measurement replicate in turn. The R package *lbfgs* was used for numerical optimization.

For datasets that sampled only two amino acids per site, we estimated RFA effects by using Fourier analysis and inferring the RFA terms from the Fourier coefficients. In a binary state space, there are fewer Fourier coefficients to model than there are reference-free effects, and the two sets of terms are easily interconvertible (Supplementary Text). The best-fit Fourier coefficients and link function were determined by the cross-validation procedure described above.

For incorporating nonspecific epistasis into reference-based analysis (RBA), the regression approach used for RFA should not be used, because regression misestimates the terms of the RBA model (Supplementary Text). For each candidate set of link function parameters, reference-based effects were computed to recapitulate the observed phenotype for mutants up to the model order. For example, the first-order model was constrained to be exact for the wild-type and its point mutants, consistent with the definition of first-order RBA. The effects and the link function were then used to predict the phenotypes of higher-order mutants, and this procedure was repeated for other parameter values to identify the link function that maximizes the *R*^2^ for higher-order mutants.

Background-averaged analysis was originally developed only for binary state space^2,25^. We extended the recursive matrix formalism to multiple states, and implemented it in a custom R script. The same multi-state formalism was recently independently derived^34^.

### Combinatorial mutagenesis datasets

We systematically mined the literature for mutagenesis experiments with a combinatorially complete design. Among the many datasets comprising fewer than 100 genotypes, we chose three datasets where high-order epistasis has been reported. Any larger dataset in which precise measurement (*r*^2^ > 0.9 between replicates) is available for at least 40% of possible genotypes was included for analysis. Several datasets were edited as described below.

The methyl-parathion hydrolase activity^43^ was measured in the presence of seven different metal cofactors. In every case, the second-order RFA with the sigmoid link function explained more than 90% of phenotypic variance. Only the Ni^2+^ dataset, in which epistasis accounts for the greatest fraction of phenotypic variance, is presented here.

The original dihydrofolate reductase dataset^3^ includes a noncoding mutation for a total of 96 variants. We only analyzed the 48 protein variants fixed for the mutant state in the noncoding site. IC75—the antibiotics concentration that reduces the growth rate by 75%—was reported in logarithmic scale, set arbitrarily as –2 when the variant is unviable at any concentration. We reverted the logarithm, making IC75 equal to 0 when the variant is unviable.

The influenza A H3N2 hemagglutinin dataset^39^ characterized an identical set of genetic variants in six different genetic backgrounds. We analyzed only the genetic background for which the measurement is most precise (Bei89).

In the avGFP dataset^13^, fluorescence is systematically higher in the second measurement replicate by a factor of 1.31. This difference was normalized when combining the two replicates.

The ParB study^44^ measures how the transcription factor ParB binds to two DNA motifs, *parS* and *NBS*. Because measurement *r*^2^ is less than 0.9 for the NBS dataset, only the *parS* dataset was analyzed. The absolute fitness of each variant was inferred by comparing the read count before and after the bulk competition assay. Variants with the pre-competition read count fewer than 15 were excluded, resulting in 42.2% coverage of the 160,000 possible genotypes—down from 97.0% in the original study.

The extent of measurement noise in the protein G GB1 domain dataset^10^ could not be directly determined because measurement was not replicated, but comparison to an independent dataset for a subset of variants showed that *r*^2^ is greater than 0.9. Variants with a pre-competition read count fewer than 100 were excluded, resulting in 68.6% coverage of the 160,000 possible genotypes—down from 93.4% in the original study.

## Acknowledgments

We thank members of the Thornton Laboratory and R. Ranganathan for discussion, and the University of Chicago Research Computing Center for high-performance computing. We thank T. Dupic, A. Phillips, and M. Desai for helpful comments on the manuscript. This work was supported by the National Institutes of Health grants R01GM131128 (J.W.T.), R01GM121931 (J.W.T.), and F32GM122251 (B.P.H.M.) and Samsung Scholarship (Y.P.).

## Author Contributions

Y.P., B.P.H.M., and J.W.T. designed research; Y.P. developed methods and analyzed data; Y.P. and J.W.T. wrote the paper.

---

**Extended Data Fig. 1.**
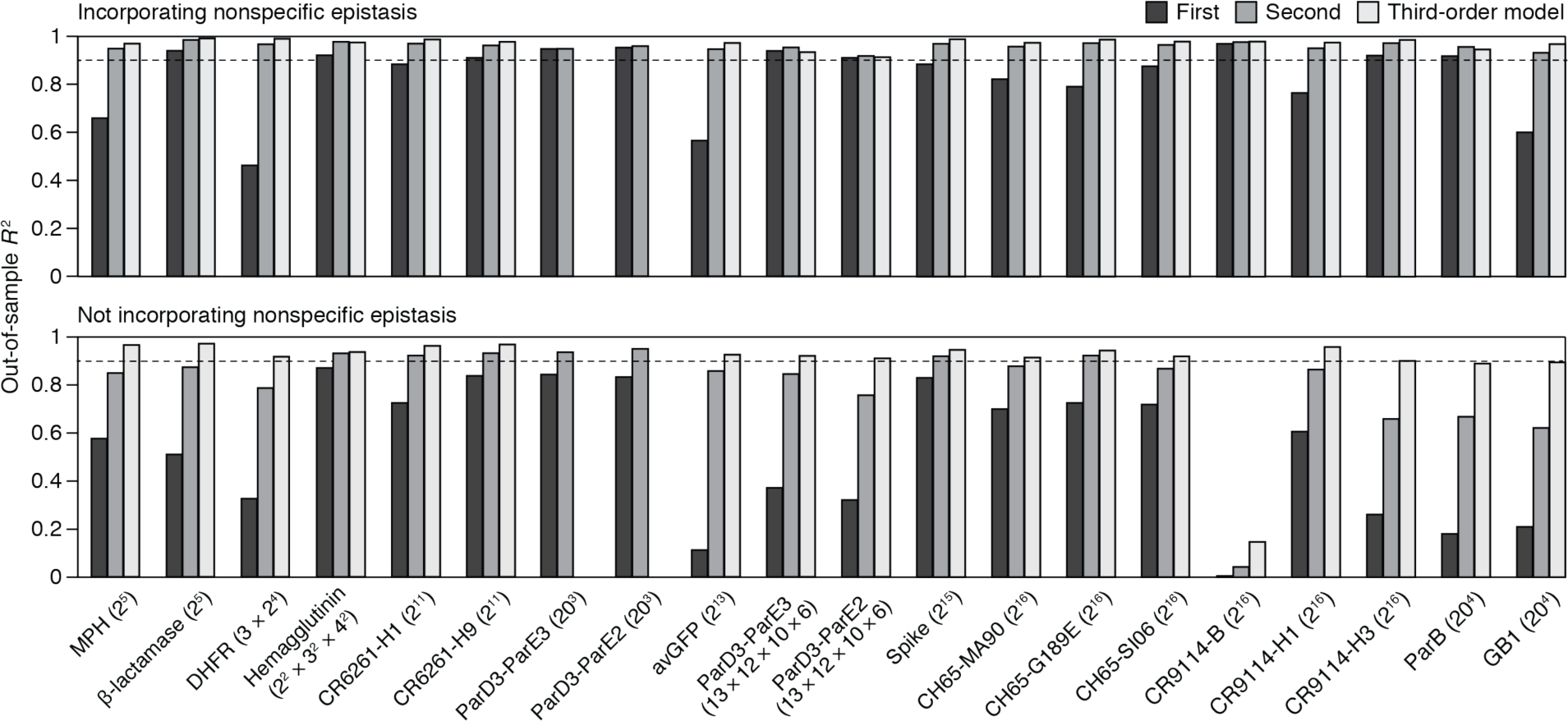
Reference-free analysis of the 20 combinatorial mutagenesis datasets. Out-of-sample *R*^2^ was computed as described in Fig. 2a with or without modeling nonspecific epistasis. Datasets are ordered by the total number of genotypes indicated in parentheses.

**Extended Data Fig. 2.**
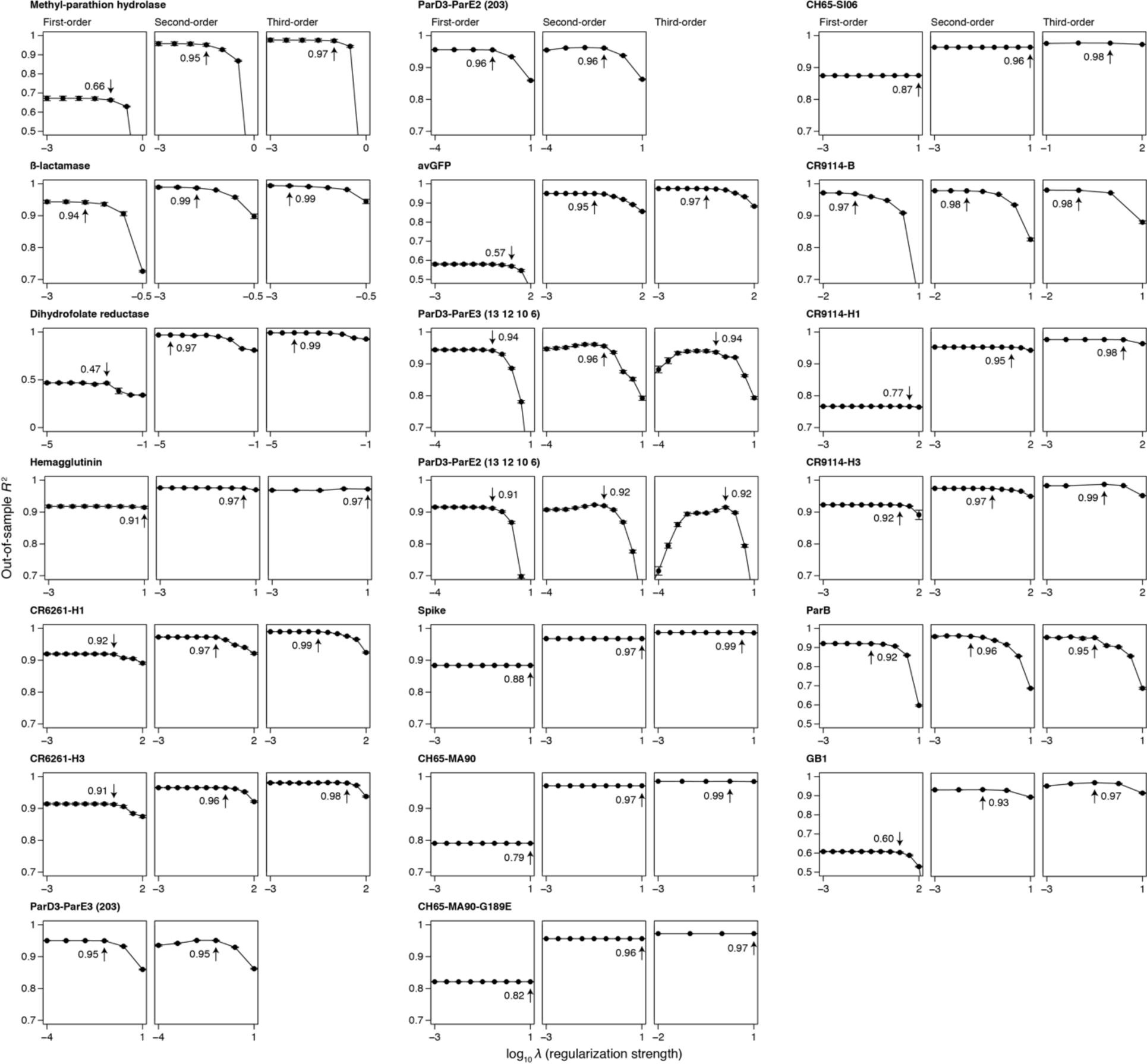
Regularization has minimal impact on the estimated variance attributed to each model order. Each panel shows the out-of-sample *R*^2^ computed under a range of L1 regularization strengths for the indicated dataset and model order. Within the panel, the dots and error bars show the mean and standard error of *R*^2^ across cross-validation replicates. The strongest regularization strength with an *R*^2^ that does not significantly differ from the maximum *R*^2^ at *p* = 0.1 was chosen. The chosen strength and the *R*^2^ at that value are marked.

