## Supporting Information for "The simplicity of protein sequence-function relationships"

### TABLE OF CONTENTS


***Comparison with other formalisms*** ... ***1***

1. ***Reference-based analysis*** ... **1**
2. ***Fourier analysis*** ... **4**
3. ***Background-averaged analysis*** ... **6**

***Exposition of reference-free analysis*** ... ***9***

1. ***Introduction*** ... **9**
2. ***Notations*** ... **9**
3. ***Definitions and interpretations*** ... **9**
4. ***Zero-mean property*** ... **11**
5. ***Generalized linear decomposition*** ... **12**
6. ***Optimal linear decomposition*** ... **12**
7. ***Variance partition*** ... **12**
8. ***Robustness to measurement noise*** ... **13**
9. ***Orthogonality of orders*** ... **13**
10. ***Unbiased estimation by regression*** ... **14**

***Appendix 1. Zero-mean property*** ... **16**

***Appendix 2. Generalized linear decomposition*** ... **17**

***Appendix 3. Optimal linear decomposition*** ... **20**

***Appendix 4. Variance partition*** ... **23**

***Appendix 6. Unbiased estimation by regression*** ... **28**

### Comparison with other formalisms

Here we provide numerical examples to contrast reference-free analysis (RFA) with reference-based, Fourier, and background-averaged analyses.

#### *1. Reference-based analysis*

*1.1. The apparent complexity of the genetic architecture depends on the choice of reference genotype*

Reference-based analysis designates a single genotype as wild-type and all others as mutants of increasing order. The phenotype is decomposed into the effects and interactions of mutations: the phenotype of a point mutant differs from the wild-type by the effect of the point mutation, that of a double mutant by the effect of the two point mutations and their pairwise interaction, and so on. The apparent complexity of the genetic architecture—the fraction of phenotypic variance attributable to each order of effects—varies depending on the choice of reference genotype. By contrast, RFA offers a unique description that minimizes the variance attributable to higher-order effects.

To illustrate this contrast, we simulated a genetic architecture consisting of 6 sites and 5 states (labeled 1 to 5) by taking the genotype (1, 1, 1, 1, 1, 1) as reference; 50, 25, and 12.5% of first-, second-, and third-order effects were sampled from the standard normal distribution, and all other effects were set to zero. We refer to the sampled genetic architecture as G1. When the phenotype is computed from the first-order effects, the squared correlation with the true phenotype is 0.30. This is the fraction of phenotypic variance explained by the first-order reference-based model defined with respect to the genotype (1, 1, 1, 1, 1, 1). The second-order model explains 78% of variance, and the third-order model 100%.

How does the same genetic architecture appear from the point of view of another genotype? Table S1 summarizes the variance partition under five randomly chosen reference genotypes.

**Table S1.** Reference-based analysis of the simulated genetic architecture G1.

| **Reference** | $R^2$**, first-order model** | $R^2$**, second-order model** | $R^2$**, third-order model** |
|---|---|---|---|
| (1, 1, 1, 1, 1, 1) | 0.30 | 0.78 | 1.00 |
| (2, 1, 3, 3, 4, 2) | 0.23 | 0.54 | 1.00 |
| (2, 5, 1, 2, 4, 1) | 0.18 | 0.65 | 1.00 |
| (1, 2, 4, 1, 4, 1) | 0.04 | 0.69 | 1.00 |
| (4, 2, 3, 1, 5, 5) | 0.06 | 0.62 | 1.00 |
| (2, 3, 3, 2, 2, 5) | 0.24 | 0.34 | 1.00 |

All analyses agree that there are up to third-order effects, but they differ markedly on the complexity of the genetic architecture. The reference genotype used for simulation offers the simplest description; this is because epistatic interactions were simulated to be sparse with respect to that genotype but this sparsity is not conserved when the reference is switched. For example, with respect to the genotype (2, 3, 3, 2, 2, 5), 90 and 45% of second- and third-order effects have a magnitude greater than 0.1. This dependence on the choice of reference genotype appropriately reflects the purpose of reference-based analysis—to understand the behavior of mutations as they accumulate on a particular genetic background.

RFA of the same genetic architecture reports $R^2$ of 0.53, 0.88 and 1.00 for the first-, second-, and third-order model, which is greater than the $R^2$ of any reference-based model of the same order.

*1.2. Sensitivity to measurement noise and missing genotypes*

A reference-based effect of order $k$ is the sum or subtraction of $2^k$ phenotypes. When individual phenotypes are measured with an error of variance ω, the variance associated with the effect is $2^k\omega$. This is always greater than ω and rises exponentially with order such that discerning effect from error in practice is possible only for the lowest few orders. By contrast, the variance associated with reference-free effects is smaller than ω for any order.

Reference-based analysis is also sensitive to missing genotypes. The reference-based effect of a mutation is defined solely by the wild-type genotype and the one mutant that differs from the wild-type by that mutation. If any of the two genotypes is missing from data, neither the mutation's effect nor any of the epistatic interactions involving that mutation can be computed. The exact computation of reference-free effects is also only possible with complete data, but because they are averages of many phenotypic measurements, they can be estimated from incomplete data by regression.

#### *1.3. Reference-based effects cannot be reliably inferred by regression*

Directly computing a mutation's effect is impossible when the mutant is missing, but the mutation also appears in other genotypes along with other mutations, so it may seem possible to disentangle its effect from those of others using regression. However, reference-based effects cannot be reliably inferred by regression. We illustrate this on the genetic architecture G1. We begin with an ideal case when every phenotype is measured precisely and therefore any effect can be computed exactly. When the first-order model defined relative to the genotype (1, 1, 1, 1, 1, 1) is fit to this dataset, the regression estimates are dramatically incorrect (Fig. S1). Furthermore, while the true model explains 30% of phenotypic variance, the inferred model explains 53%. When the second-order model is fit, the effects are estimated more accurately but still not exactly; the fraction of variance explained is also overestimated as 88% instead of 78%. The estimates are exact only with the third-order model—when all the mutational effects and interactions in the genetic architecture are modeled.

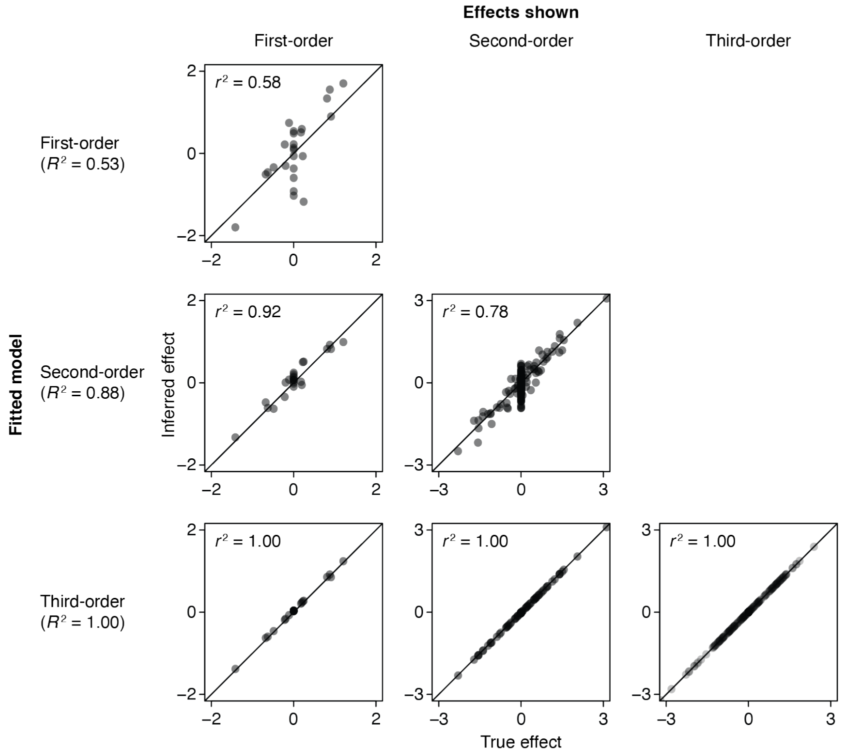

**Fig. S1 | Reference-based effects cannot be reliably estimated by regression.** Reference-based models defined with respect to the genotype (1, 1, 1, 1, 1, 1) were fit to the genetic architecture G1 in the absence of measurement noise and missing genotypes. Each panel compares the effects of a specified order with their estimates obtained using the model of indicated order.

Why does regression yield incorrect estimates of reference-based effects and overestimate their variance contribution even in the absence of measurement noise and missing genotypes? Regression finds the parameter values that minimize the sum of squared errors and therefore maximize the variance explained, but reference-based effects hold no such relation with phenotypic variance. When higher-order effects shape the phenotype but are not represented in the regression model, regression optimizes the lower-order effects to incorporate the deviations caused by the higher-order effects. Consider a simple case of two sites and two states. Suppose that genotypes (1, 1), (1, 2), and (2, 1) have a phenotype of 0 and (2, 2) a phenotype of 1. With (1, 1) as wild-type, the zero-order term $r_0 = 0$, the two additive effects $r_1 = r_2 = 0$, and the pairwise interaction $r_{12} = 1$. The first-order model predicts a phenotype of 0 for all genotypes and therefore does not explain any phenotypic variance. However, fitting this model by regression gives $r_0 = -0.25$ and $r_1 = r_2 = 0.5$ because these values minimize the sum of squared errors, explaining 67% of phenotypic variance.

Regression does not suffer from these issues when coupled with RFA. In the absence of measurement noise and missing genotypes, the inferred effects and variance contribution are exact (Fig. S2). This correspondence arises because reference-free effects minimize the sum of squared errors of phenotypic prediction.

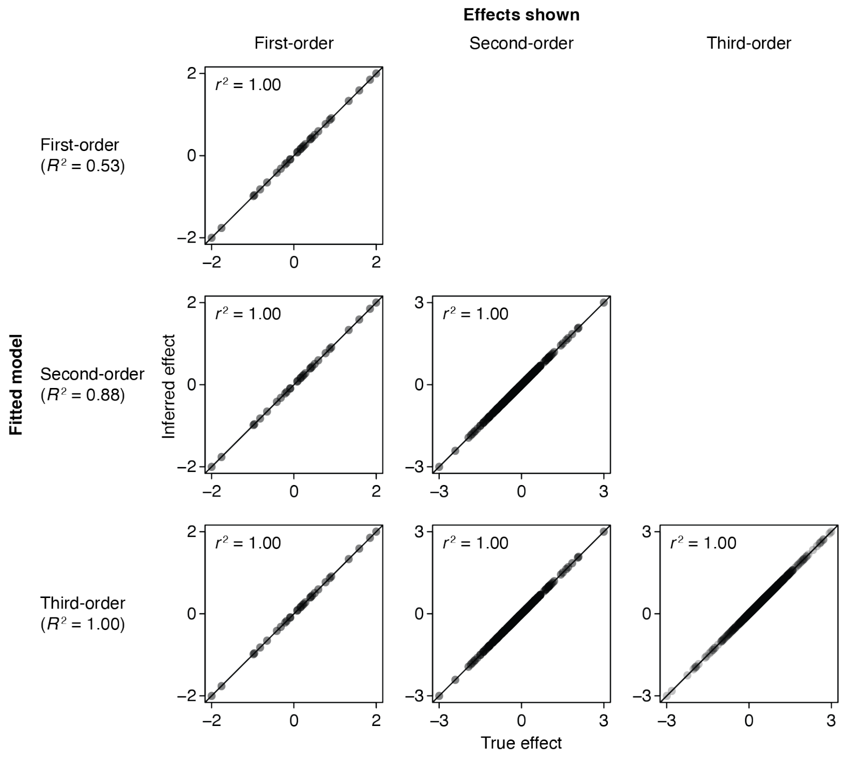


**Fig. S2 | Regression reliably estimates reference-free effects.** Reference-free models were fit to the genetic architecture G1 in the absence of measurement noise and missing genotypes. Each panel compares the effects of a specified order with their estimates obtained using the model of indicated order.

Note that reference-based models fit by regression explain the same fraction of phenotypic variance as reference-free models (53, 88, and 100% at the first, second, and third order). The two models in fact predict the same phenotype for any given genotype, because that is the prediction that minimizes the sum of squared errors. This equivalence arises because the two models have the same degrees of freedom and therefore fit the data with the same degree of accuracy. As shown above, however, the accuracy of the inferred effects contrasts strikingly between the two models. While reference-based analysis may be coupled with regression for the sole purpose of phenotypic prediction, interpreting the inferred effects as reference-based effects would mischaracterize the genetic architecture.

In Fig. S1, the regression estimates of reference-based effects are exact for the third-order model, but this is only because measurement noise and missing genotypes were not simulated. The estimates are highly inaccurate when both are present (Fig. S3). Reference-free effects, by contrast, are inferred robustly.

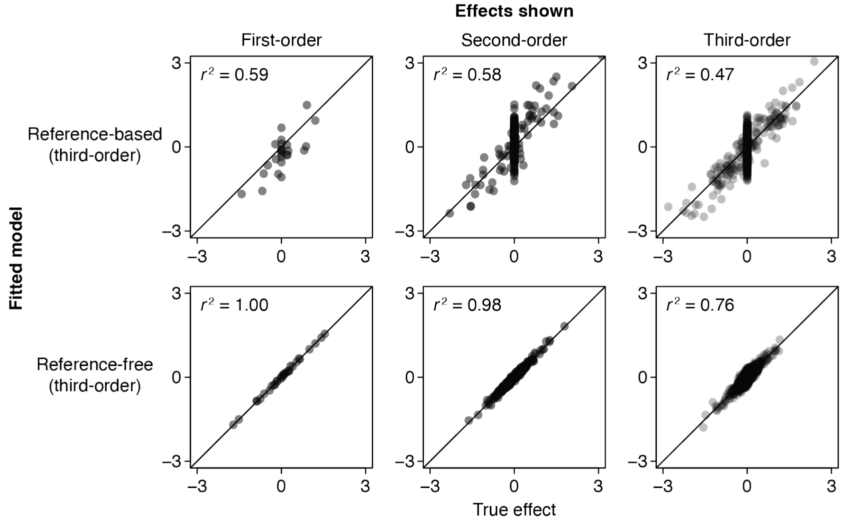


**Fig. S3 | Regression estimates of reference-based effects are sensitive to measurement noise and missing genotypes.** Measurement noise was simulated on the genetic architecture G1 to account for 5% of phenotypic variance, and the third-order reference-based or reference-free model was fit to 20% of randomly sampled genotypes. Each panel compares the effects of a specified order with their estimates.

In summary, the goal of reference-based analysis is to characterize the effects and interactions of mutations on a particular genetic background, whereas that of regression is to minimize the sum of squared errors of phenotypic prediction. Because of this misfit, the regression estimates of reference-based effects can be strikingly wrong even when obtained from precise and complete data. Avoiding this problem requires modeling all potential mutational effects and interactions in the genetic architecture. Even so, the regression estimates are sensitive to measurement noise and missing genotypes.

### 2. *Fourier analysis*

#### *2.1. The model terms (Fourier coefficients) lack straightforward genetic or biochemical meaning*

Whereas RFA decomposes the phenotype into the global effects and interactions of states, Fourier analysis decomposes the phenotype into abstract quantities called Fourier coefficients. The definition of Fourier coefficients, their relation to the phenotype, and their possible interpretation are best illustrated by an example. Consider a two-site, four-state DNA sequence, which has 16 possible genotypes. There are 16 Fourier coefficients: one zero-order coefficient ($f_0$), three first-order coefficients at each site (prefixed

$f_1$ and $f_2$), and nine second-order coefficients (prefixed $f_{1,2}$). The 16 coefficients map to the 16 phenotypes as:

$$\begin{bmatrix} y_{AA} \\ y_{AC} \\ y_{AG} \\ y_{AT} \\ y_{CA} \\ y_{CC} \\ y_{CG} \\ y_{CT} \\ y_{GA} \\ y_{GC} \\ y_{GG} \\ y_{GT} \\ y_{TA} \\ y_{TC} \\ y_{TG} \\ y_{TT} \end{bmatrix} = \begin{bmatrix} 1 & 1 & -1 & -1 & 1 & -1 & -1 & 1 & -1 & -1 & -1 & 1 & 1 & -1 & 1 & 1 \\ 1 & 1 & -1 & -1 & -1 & 1 & -1 & -1 & 1 & -1 & 1 & -1 & 1 & 1 & -1 & 1 \\ 1 & 1 & -1 & -1 & -1 & -1 & 1 & -1 & -1 & 1 & 1 & 1 & -1 & 1 & 1 & -1 \\ 1 & 1 & -1 & -1 & 1 & 1 & 1 & 1 & 1 & 1 & -1 & -1 & -1 & -1 & -1 & -1 \\ 1 & -1 & 1 & -1 & 1 & -1 & -1 & -1 & 1 & 1 & 1 & -1 & -1 & -1 & 1 & 1 \\ 1 & -1 & 1 & -1 & -1 & 1 & -1 & 1 & -1 & 1 & -1 & 1 & -1 & 1 & -1 & 1 \\ 1 & -1 & 1 & -1 & -1 & -1 & 1 & 1 & 1 & -1 & -1 & -1 & 1 & 1 & 1 & -1 \\ 1 & -1 & 1 & -1 & 1 & 1 & 1 & -1 & -1 & -1 & 1 & 1 & 1 & -1 & -1 & -1 \\ 1 & -1 & -1 & 1 & 1 & -1 & -1 & -1 & 1 & 1 & -1 & 1 & 1 & 1 & -1 & -1 \\ 1 & -1 & -1 & 1 & -1 & 1 & -1 & 1 & -1 & 1 & 1 & -1 & 1 & -1 & 1 & -1 \\ 1 & -1 & -1 & 1 & -1 & -1 & 1 & 1 & 1 & -1 & 1 & 1 & -1 & -1 & -1 & 1 \\ 1 & -1 & -1 & 1 & 1 & 1 & 1 & -1 & -1 & -1 & -1 & -1 & -1 & 1 & 1 & 1 \\ 1 & 1 & 1 & 1 & 1 & -1 & -1 & 1 & -1 & -1 & 1 & -1 & -1 & 1 & -1 & -1 \\ 1 & 1 & 1 & 1 & -1 & 1 & -1 & -1 & 1 & -1 & -1 & 1 & -1 & -1 & 1 & -1 \\ 1 & 1 & 1 & 1 & -1 & -1 & 1 & -1 & -1 & 1 & -1 & -1 & 1 & -1 & -1 & 1 \\ 1 & 1 & 1 & 1 & 1 & 1 & 1 & 1 & 1 & 1 & 1 & 1 & 1 & 1 & 1 & 1 \end{bmatrix} \begin{bmatrix} f_0 \\ f_1(1) \\ f_1(2) \\ f_1(3) \\ f_2(1) \\ f_2(2) \\ f_2(3) \\ f_{1,2}(1,1) \\ f_{1,2}(1,2) \\ f_{1,2}(1,3) \\ f_{1,2}(2,1) \\ f_{1,2}(2,2) \\ f_{1,2}(2,3) \\ f_{1,2}(3,1) \\ f_{1,2}(3,2) \\ f_{1,2}(3,3) \end{bmatrix}.$$

The phenotype of any genotype is a signed sum of all 16 Fourier coefficients, with different sets of signs distinguishing different genotypes. This contrasts with the simplicity of RFA, where each phenotype is a simple sum of only four terms—the intercept, two additive effects, and one pairwise interaction.

Interpreting Fourier coefficients based on this mapping is not straightforward. One way to understand Fourier coefficients is to relate them to reference-free effects. With some algebra, the first-order Fourier coefficients at site 1 can be written as,

$$f_1(1) = \frac{e_1(A)+e_1(T)}{2},$$
$$f_1(2) = \frac{e_1(C)+e_1(T)}{2},$$
$$f_1(3) = \frac{e_1(G)+e_1(T)}{2}.$$

That is, $f_1(1)$ is the average reference-free effect of A and T, or of the composite state W. Similarly, $f_1(2)$ and $f_1(3)$ are each the reference-free effect of the composite state Y or K, respectively. This relatively simple interpretation is possible because DNA has just four states. For proteins, each of the 19 first-order Fourier coefficients is the average of the reference-free effects of a unique set of 10 amino acids, for which there would be no biochemically or genetically interpretable meaning.

The relationship between Fourier coefficients and reference-free effects is much more complicated beyond the first order, precluding any straightforward interpretation for higher-order Fourier coefficients. For example,

$$f_{1,2}(1,1) = \frac{1}{4}\left[\frac{e_{1,2}(A,A) - e_{1,2}(A,C) - e_{1,2}(A,G) + e_{1,2}(A,T)}{4} + \frac{e_{1,2}(T,A) - e_{1,2}(T,C) - e_{1,2}(T,G) + e_{1,2}(T,T)}{4} + \frac{e_{1,2}(A,A) - e_{1,2}(C,A) - e_{1,2}(G,A) + e_{1,2}(T,A)}{4} + \frac{e_{1,2}(A,T) - e_{1,2}(C,T) - e_{1,2}(G,T) + e_{1,2}(T,T)}{4}\right].$$

Roughly, this is the average pairwise interaction between (A or T) at one site and (A or T) at the other site minus the average pairwise interaction between (A or T) at one site and (C or G) at the other. With 20 states, the pairwise Fourier terms are accordingly more elaborate.

*3.2. Fourier analysis is a compact encoding of RFA*

Fourier analysis of the two-nucleotide DNA involved 16 coefficients, equal in number to the 16 possible genotypes. RFA of the same genetic architecture requires 25 terms—the intercept, four first-order effects for each site, and 16 pairwise interactions. In general, with $n$ sites and $q$ states, there are $nq$ first-order reference-free effects, $q$ for each site. There are $\binom{n}{2}q^2$ second-order effects, $q^2$ for each pair of sites. With $\binom{n}{k}q^k$ effects of order $k$, the total number of reference-free effects is $(q + 1)^n$. By contrast, the number of Fourier coefficients of order $k$ is $\binom{n}{k}(q-1)^k$. The total number is $q^n$, which is the minimal necessary number to describe the $q^n$ possible genotypes.

Fourier analysis can be considered a compact encoding of RFA, where the $q$ states are encoded using $(q - 1)$ Fourier bases. For DNA, the three Fourier bases correspond to the three familiar nucleotide groupings: $w$ ("weak" nucleotides A and T vs. "strong" nucleotides C and G), $y$ (pyrimidines C and T vs. purines A and G), and $k$ (keto-containing nucleotides G and T vs. non-keto nucleotides A and C). A is encoded in the *wyk*-space as (1, –1, –1). The phenotype of the sequence A is decomposed as $f_0 + f_1(\text{W}) - f_1(\text{Y}) - f_1(\text{K})$, and therefore the reference-free effect $e_1(\text{A})$ equals $f_1(\text{W}) - f_1(\text{Y}) - f_1(\text{K})$. Fourier coefficients and reference-free effects can be interconverted through such linear mappings. Moreover, in the absence of missing genotypes, RFA and Fourier analysis of the same order predict the same phenotype for any given genotype.

The compact encoding of Fourier analysis has benefits and costs. The numerical economy compared with RFA is substantial at high orders, especially when the number of states is small. With two states, there are $2^{10} = 1{,}024$ tenth-order reference-free effects but just one Fourier coefficient for each set of ten sites. The degree of compression is minor for proteins, however, where the number of Fourier coefficients is 19 for a site as opposed to 20, and 361 for a site-pair as opposed to 400. Moreover, the numerical economy comes at the cost of the aforementioned lack of straightforward interpretation, the convoluted mapping from coefficients to phenotype, and extra sensitivity to missing genotypes (Fig. 1e). In practice, Fourier analysis may be preferrable over RFA for binary state spaces where the numerical economy relative to RFA is maximal, the coefficients have a simple relation to reference-free effects, and the extra sensitivity to missing genotypes is small.

### *3. Background-averaged analysis*

Background-averaged analysis differs from RFA in that it decomposes the phenotype into the effects of mutations, not states. It also differs from reference-based analysis because it examines the average effects of mutations across all genetic backgrounds, rather than their effects on one particular background. We illustrate background-averaged analysis and explain why it is more sensitive to measurement noise and missing genotypes than is RFA.

We consider a genotype space of two sites and three states (0, 1, and 2). Background-averaged analysis requires a choice of wild-type state for each site. We choose state 0 as wild-type and calculate the effects of mutations to states 1 and 2. The intercept is defined as the mean phenotype of all genotypes:

$$b_0 = \bar{y}.$$

The first-order effect of state 1 in site 1, denoted by $b_{1\cdot}$, is the effect of mutating state 0 to 1 measured on the background of each possible state in site 2:

$$b_{1\cdot} = \frac{(y_{10}-y_{00})+(y_{11}-y_{01})+(y_{12}-y_{02})}{3}.$$

Likewise, the first-order effect of state 1 in site 2, denoted by $b_{\cdot 1}$, is the effect of mutating state 0 to 1 measured on the background of each possible state in site 1:

$$b_{\cdot 1} = \frac{(y_{01} - y_{00}) + (y_{11} - y_{10}) + (y_{21} - y_{20})}{3}.$$

Because there are only two sites in this example, there is no genetic background over which to average pairwise interactions. Therefore, background-averaged interactions are identical to the corresponding reference-based interactions. For example,

$$b_{11} = y_{11} - y_{10} - y_{01} + y_{00}.$$

These definitions can be stated compactly as matrix multiplication:

$$\begin{bmatrix} b_0 \\ b_{\cdot 1} \\ b_{\cdot 2} \\ b_{1\cdot} \\ b_{11} \\ b_{12} \\ b_{2\cdot} \\ b_{21} \\ b_{22} \end{bmatrix} = \begin{bmatrix} \frac{1}{9} & \frac{1}{9} & \frac{1}{9} & \frac{1}{9} & \frac{1}{9} & \frac{1}{9} & \frac{1}{9} & \frac{1}{9} & \frac{1}{9} \\ -\frac{1}{3} & \frac{1}{3} & 0 & -\frac{1}{3} & \frac{1}{3} & 0 & -\frac{1}{3} & \frac{1}{3} & 0 \\ -\frac{1}{3} & 0 & \frac{1}{3} & -\frac{1}{3} & 0 & \frac{1}{3} & -\frac{1}{3} & 0 & \frac{1}{3} \\ -\frac{1}{3} & -\frac{1}{3} & -\frac{1}{3} & \frac{1}{3} & \frac{1}{3} & \frac{1}{3} & 0 & 0 & 0 \\ 1 & -1 & 0 & -1 & 1 & 0 & 0 & 0 & 0 \\ 1 & 0 & -1 & -1 & 0 & 1 & 0 & 0 & 0 \\ -\frac{1}{3} & -\frac{1}{3} & -\frac{1}{3} & 0 & 0 & 0 & \frac{1}{3} & \frac{1}{3} & \frac{1}{3} \\ 1 & -1 & 0 & 0 & 0 & 0 & -1 & 1 & 0 \\ 1 & 0 & -1 & 0 & 0 & 0 & -1 & 0 & 1 \end{bmatrix} \begin{bmatrix} y_{00} \\ y_{01} \\ y_{02} \\ y_{10} \\ y_{11} \\ y_{12} \\ y_{20} \\ y_{21} \\ y_{22} \end{bmatrix}.$$

The structure of this matrix explains why background-averaged analysis is more sensitive to measurement noise and missing genotypes than is RFA. To see this, we represent RFA as matrix multiplication, showing only state 1 for simplicity:

$$\begin{bmatrix} e_0 \\ e_1(0) \\ e_1(1) \\ e_2(0) \\ e_2(1) \\ e_{1,2}(0,0) \\ e_{1,2}(0,1) \\ e_{1,2}(1,0) \\ e_{1,2}(1,1) \end{bmatrix} = \begin{bmatrix} \frac{1}{9} & \frac{1}{9} & \frac{1}{9} & \frac{1}{9} & \frac{1}{9} & \frac{1}{9} & \frac{1}{9} & \frac{1}{9} & \frac{1}{9} \\ \frac{2}{9} & \frac{2}{9} & \frac{2}{9} & -\frac{1}{9} & -\frac{1}{9} & -\frac{1}{9} & -\frac{1}{9} & -\frac{1}{9} & -\frac{1}{9} \\ -\frac{1}{9} & -\frac{1}{9} & -\frac{1}{9} & \frac{2}{9} & \frac{2}{9} & \frac{2}{9} & -\frac{1}{9} & -\frac{1}{9} & -\frac{1}{9} \\ \frac{2}{9} & -\frac{1}{9} & -\frac{1}{9} & \frac{2}{9} & -\frac{1}{9} & -\frac{1}{9} & \frac{2}{9} & -\frac{1}{9} & -\frac{1}{9} \\ -\frac{1}{9} & \frac{2}{9} & -\frac{1}{9} & -\frac{1}{9} & \frac{2}{9} & -\frac{1}{9} & -\frac{1}{9} & \frac{2}{9} & -\frac{1}{9} \\ \frac{4}{9} & -\frac{2}{9} & -\frac{2}{9} & -\frac{2}{9} & \frac{1}{9} & \frac{1}{9} & -\frac{2}{9} & \frac{1}{9} & \frac{1}{9} \\ -\frac{2}{9} & \frac{4}{9} & -\frac{2}{9} & \frac{1}{9} & -\frac{2}{9} & \frac{1}{9} & \frac{1}{9} & -\frac{2}{9} & \frac{1}{9} \\ -\frac{2}{9} & \frac{1}{9} & \frac{1}{9} & \frac{4}{9} & -\frac{2}{9} & -\frac{2}{9} & -\frac{2}{9} & \frac{1}{9} & \frac{1}{9} \\ \frac{1}{9} & -\frac{2}{9} & \frac{1}{9} & -\frac{2}{9} & \frac{4}{9} & -\frac{2}{9} & \frac{1}{9} & -\frac{2}{9} & \frac{1}{9} \end{bmatrix} \begin{bmatrix} y_{00} \\ y_{01} \\ y_{02} \\ y_{10} \\ y_{11} \\ y_{12} \\ y_{20} \\ y_{21} \\ y_{22} \end{bmatrix}.$$

Many of the matrix elements for background-averaged analysis are 1 or –1, indicating that the noise in phenotypic measurement is directly propagated to the effects. The many zeros also mean that the effects are calculated by averaging over only subsets of genotypes. By contrast, all the matrix elements for RFA are nonzero and less than 4/9 in magnitude.

In background-averaged analysis, $y_{00}$ contributes to all nine effects whereas $y_{11}$ contributes to only four. Genotypes containing the wild-type states contribute disproportionately to background-averaged effects, causing extra sensitivity to noise and missing data for these effects. By contrast, all genotypes contribute with equal weights to reference-free effects.

Although background-averaged effects are defined intuitively as the average effects of mutations, their mapping back to phenotype is complicated:

$$\begin{bmatrix} y_{00} \\ y_{01} \\ y_{02} \\ y_{10} \\ y_{11} \\ y_{12} \\ y_{20} \\ y_{21} \\ y_{22} \end{bmatrix} = \begin{bmatrix} 1 & -\frac{1}{3} & -\frac{1}{3} & -\frac{1}{3} & \frac{1}{9} & \frac{1}{9} & -\frac{1}{3} & \frac{1}{9} & \frac{1}{9} \\ 1 & \frac{2}{3} & -\frac{1}{3} & -\frac{1}{3} & -\frac{2}{9} & \frac{1}{9} & -\frac{1}{3} & -\frac{2}{9} & \frac{1}{9} \\ 1 & -\frac{1}{3} & \frac{2}{3} & -\frac{1}{3} & \frac{1}{9} & -\frac{2}{9} & -\frac{1}{3} & \frac{1}{9} & -\frac{2}{9} \\ 1 & -\frac{1}{3} & -\frac{1}{3} & \frac{2}{3} & -\frac{2}{9} & -\frac{2}{9} & -\frac{1}{3} & \frac{1}{9} & \frac{1}{9} \\ 1 & \frac{2}{3} & -\frac{1}{3} & \frac{2}{3} & \frac{4}{9} & -\frac{2}{9} & -\frac{1}{3} & -\frac{2}{9} & \frac{1}{9} \\ 1 & -\frac{1}{3} & \frac{2}{3} & \frac{2}{3} & -\frac{2}{9} & \frac{4}{9} & -\frac{1}{3} & \frac{1}{9} & -\frac{2}{9} \\ 1 & -\frac{1}{3} & -\frac{1}{3} & -\frac{1}{3} & \frac{1}{9} & \frac{1}{9} & \frac{2}{3} & -\frac{2}{9} & -\frac{2}{9} \\ 1 & \frac{2}{3} & -\frac{1}{3} & -\frac{1}{3} & -\frac{2}{9} & \frac{1}{9} & \frac{2}{3} & \frac{4}{9} & -\frac{2}{9} \\ 1 & -\frac{1}{3} & \frac{2}{3} & -\frac{1}{3} & \frac{1}{9} & -\frac{2}{9} & \frac{2}{3} & -\frac{2}{9} & \frac{4}{9} \end{bmatrix} \begin{bmatrix} b_0 \\ b_{\cdot 1} \\ b_{\cdot 2} \\ b_{1\cdot} \\ b_{11} \\ b_{12} \\ b_{2\cdot} \\ b_{21} \\ b_{22} \end{bmatrix}.$$

The phenotype of each genotype is a weighted sum of all background-averaged effects, including those that represent mutations not in the genotype of interest. This means that the error in the estimate of one effect propagates to all genotypes. By contrast, the mapping from reference-free effects to phenotype is sparse, so the impact of error in the estimate of an effect is limited to the small subset of genotypes involving that effect.

### Exposition of reference-free analysis

#### *1. Introduction*

Here we provide a formal exposition of reference-free analysis (RFA), including proofs for its key properties. We begin by defining and interpreting reference-free effects. We develop the notion of generalized linear decomposition, a unified formalism for representing any linear decomposition of genetic architecture, which we use to show that RFA explains the maximum fraction of phenotypic variance that can be explained by any linear decomposition of the same order. Next, we show that the phenotypic variance can be decomposed into the contribution of each reference-free effect, which enables the variance partition framework for quantifying and comparing the phenotypic contribution of any effect or set of effects. We end by showing that reference-free effects can be robustly computed from noisy phenotypic measurements and that they can be estimated from an incomplete sample of genotype space by regression.

#### *2. Notations*

RFA can be applied to any discrete-state genetic architecture, but for simplicity we consider a genotype space with the same number of states ($q$) across sites. The definitions and proofs provided here can be easily extended to genotype spaces with different numbers of states among sites. The $n$-tuple $\boldsymbol{g} = (g_1, \ldots, g_n)$ denotes a genotype with state $g_i$ in site $i = 1, \ldots, n$. The phenotype of $\boldsymbol{g}$ is written as $y(\boldsymbol{g})$. The set of all genotypes is denoted by $G$, and the set of all genotypes sharing states $s_1, \ldots, s_k$ in sites $i_1, \ldots, i_k$ is denoted by $G_{i_1,\ldots,i_k}^{s_1,\ldots,s_k}$. Angled brackets denote averaging over a set; for example, $\langle y|G \rangle$ is the average phenotype of all genotypes. The set $N = \{1, \ldots, n\}$ is used to denote iteration over sites and site-combinations; $\Sigma_{i \in N}$ indicates summation over all sites and $\Sigma_{i_1 < i_2 \in N}$ over all site-pairs. Likewise, $Q = \{1, \ldots, q\}$ is used to denote iteration over states and state-combinations.

We do not consider nonspecific epistasis here. Nonspecific epistasis can be incorporated by modeling $y(\boldsymbol{g})$ as a nonlinear transformation of the genetic score $s(\boldsymbol{g})$, which would be defined according to the formalism below.

#### *3. Definitions and interpretations*

We first present RFA as a stepwise approximation of genetic architecture. Two alternative interpretations are then presented.

The intercept or zero-order term, $e_0$, is defined as the mean phenotype of all genotypes:

$$e_0 = \langle y|G \rangle.$$

This is the best single-parameter approximation of the genetic architecture in the sense that it minimizes the mean squared error of phenotypic prediction across sequence space.

The first-order term representing state $s$ in site $i$, denoted by $e_i(s)$, is defined as

$$e_i(s) = \langle y|G_i^s \rangle - e_0.$$

It can be considered the error associated with approximating $\langle y|G_i^s \rangle$—the mean phenotype of the subspace comprising all genotypes sharing state $s$ in site $i$—by the lower-order term $e_0$ (the mean phenotype across the entire genotype space).

The above expression extends naturally to higher-order terms. The second-order term for the state-pair ($s_1$, $s_2$) in site-pair ($i_1$, $i_2$), denoted by $e_{i_1,i_2}(s_1,s_2)$, is the error associated with approximating $\left\langle y \middle| G_{i_1,i_2}^{s_1,s_2} \right\rangle$ using the lower-order terms:

$$e_{i_1,i_2}(s_1,s_2) = \left\langle y \middle| G_{i_1,i_2}^{s_1,s_2} \right\rangle - \left[e_0 + e_{i_1}(s_1) + e_{i_2}(s_2)\right].$$

In general, the $k$-th-order term for the state-combination ($s_1$, …, $s_k$) in site-combination ($i_1$, …, $i_k$), denoted by $e_{i_1,\dots,i_k}(s_1,\dots,s_k)$, is defined as

$$e_{i_1,\dots,i_k}(s_1,\dots,s_k) = \left\langle y \middle| G_{i_1,\dots,i_k}^{s_1,\dots,s_k} \right\rangle - \left\langle y \middle| G_{i_1,\dots,i_k}^{s_1,\dots,s_k} \right\rangle_{(k-1)}, \quad (1)$$

where the subscript ($k$ – 1) indicates approximation using terms of order up to ($k$ – 1):

$$\left\langle y \middle| G_{i_1,\dots,i_k}^{s_1,\dots,s_k} \right\rangle_{(k-1)} = e_0 + \sum_{\alpha \in K} e_{i_\alpha}(s_\alpha) + \sum_{\alpha_1<\alpha_2 \in K} e_{i_{\alpha_1},i_{\alpha_2}}(s_{\alpha_1},s_{\alpha_2}) + \dots + \sum_{\alpha_1<\dots<\alpha_{k-1} \in K} e_{i_{\alpha_1},\dots,i_{\alpha_{k-1}}}(s_{\alpha_1},\dots,s_{\alpha_{k-1}}),$$

where $K$ denotes the set $\{1, \dots, k\}$.

This stepwise process builds an increasingly refined approximation of genetic architecture. The zero-order term is the crudest approximation, predicting every phenotype by the mean. The first-order terms describe the mean phenotypes of $nq$ subspaces, each consisting of all genotypes that share a particular state in a particular site. The second-order terms describe the mean phenotypes of smaller subspaces, each consisting of all genotypes sharing a pair of states in a pair of sites. Higher-order terms offer finer descriptions, with the highest-order terms describing the phenotypes of the smallest subspaces—individual genotypes.

We have so far presented reference-free terms as errors associated with lower-order approximations. They can also be interpreted as phenotypic effects. For example, the first-order term $e_i(s)$ quantifies how the mean phenotype of the genotypes sharing the state $s$ in site $i$ differs from that of all genotypes; it can therefore be interpreted as the average phenotypic effect of that state. Similarly, the second-order term $e_{i_1,i_2}(s_1,s_2)$ quantifies how the mean phenotype of the genotypes sharing the state-pair ($s_1$, $s_2$) in site-pair ($i_1$, $i_2$) differs from that of all genotypes when the first-order effects are accounted for; it is the average epistatic effect of the state-pair.

A third perspective on RFA interprets the terms of order $k$ as measuring the context-dependence of terms of order ($k$ – 1). Let us re-think the definition of the first-order term

$$e_i(s) = \langle y|G_i^s\rangle - e_0.$$

$\langle y|G_i^s\rangle$ can be considered the intercept of a genotype space consisting only of genotypes with state $s$ in site $i$. We denote this relationship by $\langle y|G_i^s\rangle = e_0|_i^s$. Thus,

$$e_i(s) = e_0|_i^s - e_0.$$

In this expression, $e_i(s)$ quantifies how different the intercept is when calculated for the complete space $G$ versus the subspace $G_i^s$. A parallel exists for second-order terms:

$$
\begin{aligned}
e_{i_1,i_2}(s_1,s_2) &= \left\langle y \middle| G_{i_1,i_2}^{s_1,s_2} \right\rangle - \left[e_0 + e_{i_1}(s_1) + e_{i_2}(s_2)\right] \\
&= \left\langle y \middle| G_{i_1,i_2}^{s_1,s_2} \right\rangle - \left\langle y \middle| G_{i_1}^{s_1} \right\rangle - \left\langle y \middle| G_{i_2}^{s_2} \right\rangle + \langle y|G\rangle \\
&= \left[\left\langle y \middle| G_{i_1,i_2}^{s_1,s_2} \right\rangle - \left\langle y \middle| G_{i_1}^{s_1} \right\rangle\right] - \left[\left\langle y \middle| G_{i_2}^{s_2} \right\rangle - \langle y|G\rangle\right] \\
&= e_{i_2}(s_2)\big|_{i_1}^{s_1} - e_{i_2}(s_2),
\end{aligned}
$$

where $e_{i_2}(s_2)\big|_{i_1}^{s_1}$ is the first-order term $e_{i_2}(s_2)$ for the subspace $G_{i_1}^{S_1}$. In general,

$$
e_{i_1,\dots,i_k}(s_1,\dots,s_k) = e_{i_2,\dots,i_k}(s_2,\dots,s_k)\big|_{i_1}^{s_1} - e_{i_2,\dots,i_k}(s_2,\dots,s_k).
$$

Here site $i_1$ is chosen as the context but any other site could also be chosen.

Finally, the phenotype of a particular genotype is the sum of the reference-free effects of all constituent states. Substituting $n$ for $k$ in Eq. (1) and noting that $\left\langle y \middle| G_{i_1,\dots,i_n}^{g_1,\dots,g_n} \right\rangle$ is simply the phenotype of $\boldsymbol{g} = (g_1, \dots, g_n)$, it follows that

$$
y(\boldsymbol{g}) = e_0 + \sum_{i\in N} e_i(g_i) + \cdots + \sum_{i_1<\cdots<i_k\in N} e_{i_1,\dots,i_k}\left(g_{i_1},\dots,g_{i_k}\right) + \cdots + e_{1,\dots,n}(g_1,\dots,g_n). \tag{2}
$$

#### *4. Zero-mean property*

Reference-free terms satisfy an equality that we call the zero-mean property. This property forms the basis of all desirable properties of RFA, and can be said to define RFA in the sense described below. We simply state the property here and prove it in Appendix 1.

For first-order terms, the zero-mean property states that the mean of all $q$ terms at a site is zero:

$$
\langle e_i|Q\rangle = \frac{1}{q}\sum_{s\in Q} e_i(s) = 0,
$$

where $Q$ denotes the set of all $q$ states. This holds for any genetic architecture by virtue of the definition of reference-free terms. For conciseness, let • denote averaging across states. For example, $e_i(\cdot)$ is the average of the $q$ first-order effects in site $i$, and $e_{i_1,i_2}(\cdot,s_2)$ is the average of the $q$ second-order effects in site-pair ($i_1$, $i_2$) that share state $s_2$ in site $i_2$. The zero-mean property for first-order terms can be restated as $e_i(\cdot) = 0$. In general, for any site-combination ($i_1$, …, $i_k$), the mean of any $q$ terms that vary across a single site is zero:

$$
e_{i_1,\dots,i_k}(\cdot,s_2,\dots,s_k) = e_{i_1,\dots,i_k}(s_1,\cdot,s_3,\dots,s_k) \dots = e_{i_1,\dots,i_k}(s_1,\dots,s_{k-1},\cdot) = 0,
$$

where ($s_1$, …, $s_k$) can be any state-combination. When the $q^2$ second-order terms for a site-pair are arranged in a $q \times q$ matrix, every row and column of the matrix sums to zero. Similarly, when the $q^3$ third-order terms for a site-triplet are arranged in a $q \times q \times q$ array, every one-dimensional section (and thus every two-dimensional section and the entire array) sums to zero.

*5. Generalized linear decomposition*

RFA linearly decomposes the phenotype into the effect of each state and state-combination. There are alternative ways of linearly decomposing the phenotype, including reference-based, Fourier, and background-averaged analyses. We now show that RFA explains the maximum fraction of phenotypic variance that can be explained by any linear decomposition of the same order. Proving this requires a common formalism to express any linear decomposition of a genetic architecture and showing that RFA is the most powerful. We call this unified formalism the generalized linear decomposition.

Consider Eq. (2) without the reference-free definition of the terms. It states a general formula for linearly decomposing the phenotype. Let us count the number of terms in this decomposition. There are $nq$ first-order terms, $q$ for each site. There are $\binom{n}{2}q^2$ second-order terms—$q^2$ pairs of states for every pair of sites. With $\binom{n}{k}q^k$ terms of order $k$, the total number of terms equals $(q + 1)^n$, which is greater than the total number of genotypes—$q^n$. This means that there are many ways of linearly decomposing the same genetic architecture, distinguished by how the terms in Eq. (2) are defined. Constraining the terms to satisfy the zero-mean property yields the reference-free decomposition: the $q$ first-order terms for a site must sum to zero, reducing their degrees of freedom to $(q – 1)$; every row and column of the $q \times q$ matrix of $q^2$ second-order terms for a site-pair must sum to zero, reducing their degrees of freedom to $(q – 1)^2$. This is the sense in which the zero-mean property defines RFA.

Alternative constraints yield alternative decompositions. For example, given a genotype $\boldsymbol{r} = (r_1, \ldots, r_n)$, setting every term involving the state $r_i$ in site $i$ to zero yields the familiar reference-based decomposition with $\boldsymbol{r}$ as the reference genotype.

We call Eq. (2) the generalized linear decomposition of genetic architecture. In Appendix 2, we show that any linear decomposition of genetic architecture is a special case of Eq. (2) obtained by subjecting the terms to a set of constraints that reduces their degrees of freedom to $q^n$. The notion of generalized linear decomposition allows us to ask the following question.

*6. Optimal linear decomposition*

Among the infinitely many ways of linearly decomposing a genetic architecture, which is the most optimal? Answering this question requires defining optimality. Given a particular linear decomposition, let $y_k(\boldsymbol{g})$ denote the phenotype of $\boldsymbol{g}$ approximated by the terms of order up to $k$—the truncation of Eq. (2) removing all higher-order terms. The accuracy of this approximation can be measured by the mean squared error:

$$\epsilon_G[y_k] = \frac{1}{q^n}\sum_{\boldsymbol{g}\in G}[y(\boldsymbol{g}) - y_k(\boldsymbol{g})]^2.$$

We define an optimal linear decomposition of order $k$ as that which minimizes $\varepsilon_G$. Appendix 3 shows that RFA minimizes $\varepsilon_G$ for any $k$ for any genetic architecture: it offers the most accurate linear approximation possible at every order. Our proof does not rule out alternative decompositions that are as accurate as RFA. When computed from complete data, Fourier coefficients and reference-free effects predict the same phenotype for any genotype.

*7. Variance partition*

A key task in understanding the causal architecture of any system is to identify the relative contribution of each causal factor to the variation in dependent variable. Intuitively, causal terms with larger coefficients imply greater importance. However, magnitude cannot be the sole criterion because terms of different order affect different numbers of genotypes. A reference-free effect of order $k$ contributes to the phenotype of one in $q^k$ genotypes; given the same magnitude, a lower-order effect is more consequential than a higher-order effect because it influences more genotypes. The magnitude and order must be jointly considered.

RFA enables a variance partition framework. This makes possible such statements as "this term explains 5% of phenotypic variance" or "the first-order terms together explain 80% of phenotypic variance." The applicability of variance partition is a unique feature of RFA enabled by its zero-mean property.

The total phenotypic variance

$$Var(y|G) = \frac{1}{q^n} \sum_{\boldsymbol{g} \in G} [y(\boldsymbol{g}) - \langle y|G \rangle]^2$$

quantifies the amount of phenotypic variation caused by genetic variation. Appendix 4 shows that it can be decomposed into the contribution of each reference-free effect:

$$Var(y|G) = \sum_{e \neq e_0} \frac{e^2}{q^{O(e)}},$$

where $e$ denotes any nonzero-order effect and $O(e)$ its order. Note that $1/q^{O(e)}$ is the fraction of genotypes whose phenotype involves the effect $e$. The variance contribution of each effect is thus the square of its magnitude normalized by the fraction of genotypes it affects. This confirms our intuition that a lower-order effect makes a greater phenotypic contribution than a higher-order effect of the same magnitude.

#### *8. Robustness to measurement noise*

We consider how noise in phenotypic measurement impacts the computation of reference-free effects. An experimentally characterized genetic architecture is a superposition of the true architecture and an unstructured architecture where each phenotype is an independent draw from a noise distribution. The reference-free effects of this unstructured architecture are the errors for the true architecture.

Consider an unstructured architecture generated by sampling each phenotype independently from a noise distribution with variance ω. The zero-order effect is the average of $q^n$ independent draws, so its variance is $\omega/q^n$. This is the error associated with the measurement of the true zero-order effect. Appendix 5 continues this calculation to higher orders, showing that the variance for an effect of order $k$ is

$$\frac{(q-1)^k}{q^n} \omega.$$

This is always smaller than ω—the noise involved in individual phenotypic measurements—and typically negligible when $k$ is small. RFA is therefore highly robust to measurement noise.

#### *9. Orthogonality of orders*

We now show that each order of reference-free effects is orthogonal to all other orders. That is, altering the genetic architecture by changing one order of effects does not change the value of other orders of effects. Here we demonstrate the orthogonality of the first-order effects from all others. Let $y_{-1}(\boldsymbol{g})$ denote the phenotypic contribution of second- and higher-order terms on the genotype $\boldsymbol{g} = (g_1, \ldots, g_n)$:

$$y_{-1}(\boldsymbol{g}) = \sum_{i_1<i_2\in N} e_{i_1,i_2}(g_{i_1}, g_{i_2}) + \sum_{i_1<i_2<i_3\in N} e_{i_1,i_2,i_3}(g_{i_1}, g_{i_2}, g_{i_3}) + \cdots + e_{1,\ldots,n}(g_1, \ldots, g_n)$$

Consider averaging $y_{-1}$ across the subset of genotypes sharing state $s$ in site $j$:

$$\begin{aligned}\langle y_{-1} | G_j^s \rangle &= \frac{1}{q^{n-1}} \sum_{\boldsymbol{g}\in G_j^s} y_{-1}(\boldsymbol{g}) \\ &= \frac{1}{q^{n-1}} \sum_{\boldsymbol{g}\in G_j^s} \left[ \sum_{i_1<i_2\in N} e_{i_1,i_2}(g_{i_1}, g_{i_2}) + \sum_{i_1<i_2<i_3\in N} e_{i_1,i_2,i_3}(g_{i_1}, g_{i_2}, g_{i_3}) + \cdots \right] \\ &= 0,\end{aligned}$$

where the last equality follows from the zero-mean property. It then follows that the values of first-order effects are independent of the values of higher-order effects:

$$\begin{aligned}e_j(s) &= \langle y | G_j^s \rangle - e_0 \\ &= \langle y_1 + y_{-1} | G_j^s \rangle - e_0 \\ &= \langle y_1 | G_j^s \rangle + \langle y_{-1} | G_j^s \rangle - e_0 \\ &= \langle y_1 | G_j^s \rangle - e_0,\end{aligned}$$

where $y_1$ denotes the phenotypic contribution of zero- and first-order effects. The subtraction by $e_0$ also makes the first-order effect independent of the intercept. Overall, the pattern of phenotypic variation produced by epistatic interactions appears as noise to the first-order model.

#### *10. Unbiased estimation by regression*

The exact computation of a complete reference-free model requires having a measured phenotype for every genotype, which is generally not feasible. However, it is possible to estimate them from a random sample of genotype space. Recall that among all linear decompositions of the same order, RFA minimizes the mean squared error of phenotypic prediction across the genotype space. Regression estimates, which minimize the mean squared error across sampled genotypes, therefore converge to the true values as sample size increases.

To formalize this idea, let $y_k$ denote the generalized linear decomposition truncated after order $k$:

$$y_k(\boldsymbol{g}) = e_0 + \sum_{i\in N} e_i(g_i) + \sum_{i_1<i_2\in N} e_{i_1,i_2}(g_{i_1}, g_{i_2}) + \cdots + \sum_{i_1<\cdots<i_k\in N} e_{i_1,\ldots,i_k}(g_{i_1}, \ldots, g_{i_k}).$$

The best-fit model $\hat{y}_k$ is found by minimizing the mean squared error across the set of sampled genotypes ($G^*$):

$$\hat{y}_k = \text{argmin} \sum_{\boldsymbol{g} \in G^*} [y(\boldsymbol{g}) - y_k(\boldsymbol{g})]^2.$$

One problem with this formulation is that, because of the degeneracy of generalized linear decomposition, there are many possible $\hat{y}_k$. This can be circumvented by performing a constrained regression where the parameters are subject to the zero-mean constraint. Alternatively, Appendix 6 shows that a simpler two-step procedure gives the same answer: an unconstrained regression is performed to find any $\hat{y}_k$ and the zero-mean constraint is enforced post hoc on the inferred model. $\hat{y}_k$ so obtained is consistent (it converges to the true reference-free model as sample size increases), and it is unbiased (the expected value of each term equals the true value). The unbiasedness follows from the orthogonality of orders: the unmodeled higher-order effects appear as noise to any lower-order model and therefore do not bias the regression.

#### *Appendix 1. Zero-mean property*

The zero-mean property of reference-free effects is that for any site-combination ($i_1$, …, $i_k$), the mean of all effects that vary across a single site is zero:

$$e_{i_1,\ldots,i_k}(\cdot, s_2, \ldots, s_k) = e_{i_1,\ldots,i_k}(s_1, \cdot, s_3, \ldots, s_k) \ldots = e_{i_1,\ldots,i_k}(s_1, \ldots, s_{k-1}, \cdot) = 0,$$

where ($s_1$, …, $s_k$) can be any state-combination. The zero-mean property is a defining feature of reference-free analysis, from which all of its useful properties proved in the next Appendices follow. We prove it by mathematical induction.

Recall that $G_i^s$ is the set of all genotypes with state $s$ in site $i$. $G_i^1, G_i^2, \ldots, G_i^q$ are nonoverlapping sets whose union is $G$. Therefore, the summation $\Sigma_{\boldsymbol{g}\in G}$ is equivalent to $\Sigma_{s\in Q}\Sigma_{\boldsymbol{g}\in G_i^s}$. Then,

$$\begin{aligned} e_i(\cdot) &= \frac{1}{q}\sum_{s\in Q} e_i(s) \\ &= \frac{1}{q}\sum_{s\in Q}[\langle y|G_i^s\rangle - \langle y|G\rangle] \\ &= \frac{1}{q}\sum_{s\in Q}\langle y|G_i^s\rangle - \langle y|G\rangle \\ &= \frac{1}{q}\sum_{s\in Q}\frac{1}{q^{n-1}}\sum_{\boldsymbol{g}\in G_i^s} y(\boldsymbol{g}) - \langle y|G\rangle \\ &= \frac{1}{q^n}\sum_{s\in Q}\sum_{\boldsymbol{g}\in G_i^s} y(\boldsymbol{g}) - \langle y|G\rangle \\ &= \frac{1}{q^n}\sum_{\boldsymbol{g}\in G} y(\boldsymbol{g}) - \langle y|G\rangle \\ &= 0. \end{aligned}$$

We now show that if the zero-mean property holds for effects of order ($k$ – 1), it also holds for effects of order $k$. Recall the definition

$$e_{i_1,\ldots,i_k}(s_1, \ldots, s_k) = e_{i_2,\ldots,i_k}(s_2, \ldots, s_k)|_{i_1}^{s_1} - e_{i_2,\ldots,i_k}(s_2, \ldots, s_k).$$

By the inductive hypothesis, the mean of the two ($k$ – 1)-th-order terms on the right-hand side is zero across any site $i_2$, …, $i_k$. The mean of the $k$-th-order term on the left is therefore zero across any site $i_2$, …, $i_k$. Conditioning on a site other than $i_1$ shows that the mean across $i_1$ is also zero. This completes the mathematical induction.

*Appendix 2. Generalized linear decomposition*

Let $e_{i_1,\dots,i_k}$ be a function mapping $k$-tuples of states into real numbers. We refer to the following expression as the $k$-th-order generalized linear decomposition of genetic architecture:

$$y_k(\boldsymbol{g}) = e_0 + \sum_{i\in N} e_i(g_i) + \sum_{i_1<i_2\in N} e_{i_1,i_2}(g_{i_1}, g_{i_2}) + \cdots + \sum_{i_1<\cdots<i_k\in N} e_{i_1,\dots,i_k}\big(g_{i_1}, \dots, g_{i_k}\big). \qquad \text{(A1)}$$

We showed that both reference-free and reference-based analyses can be represented as above with suitable choices of $e$. We now argue that any linear decomposition of genetic architecture can be represented as above with suitable choices of $e$. The validity of this statement depends on what a linear decomposition of genetic architecture is. Below we define it in the broadest possible sense and show that it has a form of Eq. (A1).

In the broadest sense, a linear decomposition of order zero is any function that approximates the phenotype of every genotype by a constant. It can thus be expressed as

$$y_0(\boldsymbol{g}) = e_0.$$

What is the broadest sense in which a function $y_1(\boldsymbol{g})$ is a first-order linear decomposition of genetic architecture? $y_1(\boldsymbol{g})$ should be able to use the information that $\boldsymbol{g}$ has the state $g_i$ in site $i$ and combine that information linearly across sites to determine its phenotype. It is not allowed, however, to use any information about what combination of states is found in what combination of sites. Any such function $y_1$ can be written as

$$y_1(\boldsymbol{g}) = \sum_j \lambda_j(\boldsymbol{g}),$$

where $\lambda_j$ is a function that can distinguish whether $\boldsymbol{g}$ has a particular state in a particular site:

$$\lambda_j(\boldsymbol{g}) = \begin{cases} \alpha_j, & \boldsymbol{g} \in G_i^s \\ \beta_j, & \boldsymbol{g} \notin G_i^s \end{cases}$$

for some state $s$, some site $i$, and some real numbers $\alpha_j$ and $\beta_j$. The number of such functions that make up $y_1(\boldsymbol{g})$ is unlimited. It can be shown that for any such function $y_1$, we can find a constant $e_0$ and functions $e_i$ such that

$$y_1(\boldsymbol{g}) = e_0 + \sum_{i\in N} e_i(g_i).$$

To prove this, we first sum all functions $\lambda$ that distinguish whether $\boldsymbol{g} \in G_i^s$ for a given $s$ and $i$ and write the sum as

$$\lambda_i^s(\boldsymbol{g}) = \begin{cases} \alpha_i^s, & \boldsymbol{g} \in G_i^s \\ \beta_i^s, & \boldsymbol{g} \notin G_i^s \end{cases}.$$

$y_1$ can be written as

$$\begin{aligned} y_1(g) &= \sum_j \lambda_j(\boldsymbol{g}) \\ &= \sum_{i\in N}\sum_{s\in Q} \lambda_i^s(\boldsymbol{g}) \\ &= \sum_{i\in N}\sum_{s\in Q} [\lambda_i^s(\boldsymbol{g}) - \beta_i^s + \beta_i^s] \\ &= \sum_{i\in N}\sum_{s\in Q} [\lambda_i^s(\boldsymbol{g}) - \beta_i^s] + \sum_{i\in N}\sum_{s\in Q} \beta_i^s. \end{aligned}$$

Note that

$$\lambda_i^s(\boldsymbol{g}) - \beta_i^s = \begin{cases} \alpha_i^s - \beta_i^s, & \boldsymbol{g} \in G_i^s \\ 0, & \boldsymbol{g} \notin G_i^s \end{cases}$$

and thus

$$\sum_{s\in Q} [\lambda_i^s(\boldsymbol{g}) - \beta_i^s] = \alpha_i^{g_i} - \beta_i^{g_i}.$$

The following choice therefore completes the proof:

$$\begin{aligned} e_0 &= \sum_{i\in N}\sum_{s\in Q} \beta_i^s, \\ e_i(s) &= \alpha_i^s - \beta_i^s. \end{aligned}$$

Higher-order linear decompositions can be defined similarly. In the broadest sense, a second-order linear decomposition can use the information that $\boldsymbol{g}$ has the state $g_i$ in site $i$ and state-pair $(g_{i_1}, g_{i_2})$ in site-pair $(i_1, i_2)$, but cannot use any information about higher-order combinations. Any such function $y_2$ can be written as

$$y_2(\boldsymbol{g}) = \sum_j \lambda_j(\boldsymbol{g}) + \sum_k \mu_k(\boldsymbol{g}),$$

where $\lambda_j$ is defined as above and $\mu_k$ is a function that can distinguish whether $\boldsymbol{g}$ has a particular pair of states in a particular pair of sites:

$$\mu_k(\boldsymbol{g}) = \begin{cases} \gamma_k, & \boldsymbol{g} \in G_{i_1,i_2}^{s_1,s_2} \\ \delta_k, & \boldsymbol{g} \notin G_{i_1,i_2}^{s_1,s_2} \end{cases}$$

for some sites $i_1$ and $i_2$, some states $s_1$ and $s_2$, and some real numbers $\gamma_k$ and $\delta_k$. For any such function $y_2$, we can find a constant $e_0$ and functions $e_i$ and $e_{i_1,i_2}$ such that

$$y_2(\boldsymbol{g}) = e_0 + \sum_{i\in N} e_i(g_i) + \sum_{i_1<i_2\in N} e_{i_1,i_2}(g_{i_1}, g_{i_2}).$$

To prove this, we again sum all functions λ that distinguish whether $\boldsymbol{g} \in G_i^s$ for a given *s* and *i* and write the sum as

$$\lambda_i^s(\boldsymbol{g}) = \begin{cases} \alpha_i^s, & \boldsymbol{g} \in G_i^s \\ \beta_i^s, & \boldsymbol{g} \notin G_i^s \end{cases},$$

and similarly sum all functions μ that distinguish whether or not $\boldsymbol{g} \in G_{i_1,i_2}^{s_1,s_2}$ and write the sum as

$$\mu_{i_1,i_2}^{s_1,s_2}(\boldsymbol{g}) = \begin{cases} \gamma_{i_1,i_2}^{s_1,s_2}, & \boldsymbol{g} \in G_{i_1,i_2}^{s_1,s_2} \\ \delta_{i_1,i_2}^{s_1,s_2}, & \boldsymbol{g} \notin G_{i_1,i_2}^{s_1,s_2} \end{cases}.$$

Following a logic similar to above, we can choose

$$e_0 = \sum_{i \in N} \sum_{s \in Q} \beta_i^s + \sum_{i_1 < i_2 \in N} \sum_{s_1,s_2 \in Q} \beta_{i_1,i_2}^{s_1,s_2},$$
$$e_i(s) = \alpha_i^s - \beta_i^s,$$
$$e_{i_1,i_2}(s_1,s_2) = \gamma_{i_1,i_2}^{s_1,s_2} - \delta_{i_1,i_2}^{s_1,s_2}.$$

In general, a linear decomposition of order *k* in the broadest sense is any function that can use information about the combination of states in any set of up to *k* sites and combine that information linearly across site-combinations to determine the phenotype. A logic similar to above can show that any such function can be written as Eq. (A1).

*Appendix 3. Optimal linear decomposition*

Recall that any $k$-th-order linear decomposition of genetic architecture can be written as

$$y_k(\boldsymbol{g}) = e_0 + \sum_{i \in N} e_i(g_i) + \sum_{i_1 < i_2 \in N} e_{i_1,i_2}(g_{i_1}, g_{i_2}) + \cdots + \sum_{i_1 < \cdots < i_k \in N} e_{i_1,\ldots,i_k}\left(g_{i_1}, \ldots, g_{i_k}\right).$$

Here we show that the reference-free definition of effects minimizes the sum of squared error

$$\epsilon_G = \sum_{\boldsymbol{g} \in G} [y(\boldsymbol{g}) - y_k(\boldsymbol{g})]^2,$$

for any genetic architecture at any value of $k$. We prove this by showing that the partial derivative of $\varepsilon_G$ with respect to a term is zero when that term is defined by the reference-free definition. Using $e$ to denote a term,

$$\begin{aligned}
\frac{\partial \epsilon_G}{\partial e} &= \frac{\partial}{\partial e} \sum_{\boldsymbol{g} \in G} [y(\boldsymbol{g}) - y_k(\boldsymbol{g})]^2 \\
&= \sum_{\boldsymbol{g} \in G} \frac{\partial}{\partial e} [y(\boldsymbol{g}) - y_k(\boldsymbol{g})]^2 \\
&= \sum_{\boldsymbol{g} \in G} -2[y(\boldsymbol{g}) - y_k(\boldsymbol{g})] \frac{\partial}{\partial e} y_k(\boldsymbol{g}).
\end{aligned}$$

The derivative of $y_k(\boldsymbol{g})$ with respect to $e$ is 1 if $y_k(\boldsymbol{g})$ involves $e$ and 0 otherwise. For example, for $e = e_i(s)$, the derivative of $y_k(\boldsymbol{g})$ is 1 for all $\boldsymbol{g} \in G_i^s$ and 0 for all $\boldsymbol{g} \notin G_i^s$. For $e = e_{i_1,\ldots,i_k}(s_1, \ldots, s_k)$, let $G_e$ denote the set $G_{i_1,\ldots,s_k}^{s_1,\ldots,s_k}$. Then,

$$\frac{\partial}{\partial e} y_k(\boldsymbol{g}) = \begin{cases} 1, & \boldsymbol{g} \in G_e \\ 0, & \boldsymbol{g} \notin G_e \end{cases}.$$

The partial derivative of $\varepsilon_G$ is then

$$\begin{aligned}
\frac{\partial \epsilon_G}{\partial e} &= \sum_{\boldsymbol{g} \in G} -2[y(\boldsymbol{g}) - y_k(\boldsymbol{g})] \frac{\partial}{\partial e} y_k(\boldsymbol{g}) \\
&= -2 \sum_{\boldsymbol{g} \in G_e} [y(\boldsymbol{g}) - y_k(\boldsymbol{g})] \\
&= -2q^{n-O(e)}[\langle y|G_e \rangle - \langle y_k|G_e \rangle],
\end{aligned}$$

where $O(e)$ is the order of $e$. The partial derivative is zero when

$$\langle y|G_e \rangle = \langle y_k|G_e \rangle.$$

In other words, an optimal linear decomposition of order $k$ is that which accurately predicts the average phenotype of any subset of genotypes defined by fixing up to $k$ sites. We defined reference-free effects to achieve just that! To formally show this, we prove the following property of reference-free analysis:

**Lemma 1.** Recall that $e_{i_1,\ldots,i_k}\left(g_{i_1},\ldots,g_{i_k}\right)$ denotes the $k$-th-order effect of state-combination $\left(g_{i_1},\ldots,g_{i_k}\right)$ in site-combination ($i_1$, …, $i_k$). Due to the zero-mean property, the mean of $e_{i_1,\ldots,i_k}\left(g_{i_1},\ldots,g_{i_k}\right)$ across all genotypes is zero. We now calculate the mean across subsets of genotypes defined by fixing the states at some sites. Consider the set $G_{j_1,\ldots,j_l}^{s_{j_1},\ldots,s_{j_l}}$, which comprises all genotypes with states $s_{j_1},\ldots,s_{j_l}$ in sites $j_1$, …, $j_l$. Lemma 1 claims that

$$\left\langle e_{i_1,\ldots,i_k}\middle|G_{j_1,\ldots,j_l}^{s_{j_1},\ldots,s_{j_l}}\right\rangle = 0$$

unless ($i_1$, …, $i_k$) is a subset of ($j_1$, …, $j_l$), in which case

$$\left\langle e_{i_1,\ldots,i_k}\middle|G_{j_1,\ldots,j_l}^{s_{j_1},\ldots,s_{j_l}}\right\rangle = e_{i_1,\ldots,i_k}\left(s_{i_1},\ldots,s_{i_k}\right).$$

This follows from the zero-mean property: if any site among $i_1$, …, $i_k$ lies outside the set ($j_1$, …, $j_l$), the above summation involves summing across all $q$ states in that site.

Before proving Lemma 1, let us see how it helps proving the optimality of reference-free analysis. For any effect $e = e_{j_1,\ldots,j_l}(s_{j_1},\ldots,s_{j_l})$, where $l \le k$,

$$\begin{aligned}\langle y_k|G_e\rangle &= \frac{1}{q^{n-l}}\sum_{g\in G_{j_1,\ldots,j_l}^{s_{j_1},\ldots,s_{j_l}}} y_k(\boldsymbol{g}) \\ &= \frac{1}{q^{n-l}}\sum_{g\in G_{j_1,\ldots,j_l}^{s_{j_1},\ldots,s_{j_l}}}\left[e_0+\sum_{i\in N}e_i(g_i)+\sum_{i_1<i_2\in N}e_{i_1,i_2}(g_{i_1},g_{i_2})+\cdots\right. \\ &\qquad \left.+\sum_{i_1<\cdots<i_k\in N}e_{i_1,\ldots,i_k}\left(g_{i_1},\ldots,g_{i_k}\right)\right].\end{aligned}$$

Due to Lemma 1, the sum for any site-combination that is not a subset of ($j_1$, …, $j_l$) is zero. Therefore, using $L$ to denote the set {1, 2, …, $l$},

$$\begin{aligned}\langle y_k|G_e\rangle = e_0 &+ \sum_{\alpha\in L}e_{j_\alpha}(s_{j_\alpha}) + \sum_{\alpha_1<\alpha_2\in L}e_{j_{\alpha_1},j_{\alpha_2}}(s_{j_{\alpha_1}},s_{j_{\alpha_2}}) + \cdots + \sum_{\alpha_1<\cdots<\alpha_{l-1}\in L}e_{j_{\alpha_1},\ldots,j_{\alpha_{l-1}}}\left(s_{j_{\alpha_1}},\ldots,s_{j_{\alpha_{l-1}}}\right) \\ &+ e_{j_1,\ldots,j_l}\left(s_{j_1},\ldots,s_{j_l}\right).\end{aligned}$$

This, by the definition of reference-free decomposition, equals $\left\langle y\middle|G_{j_1,\ldots,j_l}^{s_{j_1},\ldots,s_{j_l}}\right\rangle = \langle y|G_e\rangle$:

$$\left\langle y\middle|G_{j_1,\ldots,j_l}^{s_{j_1},\ldots,s_{j_l}}\right\rangle = \left\langle y\middle|G_{j_1,\ldots,j_l}^{s_{j_1},\ldots,s_{j_l}}\right\rangle_{(l-1)} + e_{j_1,\ldots,j_l}\left(s_{j_1},\ldots,s_{j_l}\right),$$

where

$$\left\langle y \middle| G_{j_1,\ldots,j_l}^{s_{j_1},\ldots,s_{j_l}} \right\rangle_{(l-1)}$$
$$= e_0 + \sum_{\alpha \in L} e_{j_\alpha}(s_{j_\alpha}) + \sum_{\alpha_1<\alpha_2 \in L} e_{j_{\alpha_1},j_{\alpha_2}}(s_{j_{\alpha_1}}, s_{j_{\alpha_2}}) + \cdots$$
$$+ \sum_{\alpha_1<\cdots<\alpha_{l-1} \in L} e_{j_{\alpha_1},\ldots,j_{\alpha_{l-1}}}\left(s_{j_{\alpha_1}}, \ldots, s_{j_{\alpha_{l-1}}}\right).$$

**Proof of Lemma 1.**

Consider first the case when $(i_1, \ldots, i_k) \subseteq (j_1, \ldots, j_l)$. Across the set $G_{j_1,\ldots,j_l}^{s_{j_1},\ldots,s_{j_l}}$, which consists only of genotypes with states $s_{j_1}, \ldots, s_{j_l}$ in sites $j_1, \ldots, j_l$, the term $e_{i_1,\ldots,i_k}(g_{i_1}, \ldots, g_{i_k})$ is a constant, $e_{i_1,\ldots,i_k}(s_{i_1}, \ldots, s_{i_k})$. This proves the second case of the lemma.

Let us now assume that *m* sites in $(i_1, \ldots, i_k)$ are outside $(j_1, \ldots, j_l)$. We make our notation flexible so that the order of sites can be permuted: for example, $e_{i_1,i_2}(s_1, s_2) = e_{i_2,i_1}(s_2, s_1)$. We can then write

$$(i_1, \ldots, i_k) = (x_1, \ldots, x_{k-m}, y_1, \ldots, y_m),$$

where $x_1, \ldots, x_{k-m}$ are inside $(j_1, \ldots, j_l)$ and $y_1, \ldots, y_m$ are outside. $G_{j_1,\ldots,j_l}^{s_{j_1},\ldots,s_{j_l}}$ can be partitioned as

$$G_{j_1,\ldots,j_l}^{s_{j_1},\ldots,s_{j_l}} = \bigcup_{t_1,\ldots,t_m \in Q} G_{j_1,\ldots,j_l,y_1,\ldots,y_m}^{s_{j_1},\ldots,s_{j_l},t_1,\ldots,t_m}.$$

Therefore, the summation $\Sigma_{g \in G_{j_1,\ldots,j_l}^{s_{j_1},\ldots,s_{j_l}}}$ equals $\Sigma_{t_1,\ldots,t_m \in Q} \Sigma_{g \in G_{j_1,\ldots,j_l,y_1,\ldots,y_m}^{s_{j_1},\ldots,s_{j_l},t_1,\ldots,t_m}}$. It follows then

$$\left\langle e_{i_1,\ldots,i_k} \middle| G_{j_1,\ldots,j_l}^{s_{j_1},\ldots,s_{j_l}} \right\rangle = \frac{1}{q^{n-l}} \sum_{g \in G_{j_1,\ldots,j_l}^{s_{j_1},\ldots,s_{j_l}}} e_{i_1,\ldots,i_k}\left(g_{i_1}, \ldots, g_{i_k}\right)$$
$$= \frac{1}{q^{n-l}} \sum_{t_1,\ldots,t_m \in Q} \sum_{g \in G_{j_1,\ldots,j_l,y_1,\ldots,y_m}^{s_{j_1},\ldots,s_{j_l},t_1,\ldots,t_m}} e_{x_1,\ldots,x_{k-m},y_1,\ldots,y_m}\left(s_{x_1}, \ldots, s_{x_{k-m}}, t_1, \ldots, t_m\right)$$
$$= \frac{q^l}{q^{n-l}} \sum_{t_1,\ldots,t_m \in Q} e_{x_1,\ldots,x_{k-m},y_1,\ldots,y_m}\left(s_{x_1}, \ldots, s_{x_{k-m}}, t_1, \ldots, t_m\right)$$
$$= 0.$$

The last equality follows from the zero-mean property.

*Appendix 4. Variance partition*

To help derive the variance partition formula, we use $Var\left(e_{i_1,\ldots,i_k}|Q^k\right)$ to denote the variance of all $k$-th-order effects at the site-combination ($i_1$, …, $i_k$).

$$Var\left(e_{i_1,\ldots,i_k}|Q^k\right) = \frac{1}{q^k}\sum_{s_1,\ldots,s_k\in Q}\left[e_{i_1,\ldots,i_k}(s_1,\ldots,s_k) - \left\langle e_{i_1,\ldots,i_k}|Q^k\right\rangle\right]^2$$
$$= \frac{1}{q^k}\sum_{s_1,\ldots,s_k\in Q} e_{i_1,\ldots,i_k}(s_1,\ldots,s_k)^2,$$

where the last equality follows from the zero-mean property. We show that the total phenotypic variance

$$V = Var(y|G) = \frac{1}{q^n}\sum_{\boldsymbol{g}\in G}[y(\boldsymbol{g}) - \langle y|G\rangle]^2$$

can be decomposed into the variance of reference-free effects at each site-combination:

$$V = \sum_{i\in N} Var(e_i|Q) + \sum_{i_1<i_2\in N} Var\left(e_{i_1,i_2}|Q^2\right) + \cdots + \sum_{i_1<\cdots<i_k\in N} Var\left(e_{i_1,\ldots,i_k}|Q^k\right) + \cdots + Var\left(e_{1,\ldots,n}|Q^n\right).$$

This is equivalent to the variance partition formula

$$V = \sum_{e\neq e_0}\frac{e^2}{q^{O(e)}}.$$

Recall that for any genotype $\boldsymbol{g} = (g_1,\ldots,g_n)$,

$$y(\boldsymbol{g}) = e_0 + \sum_{i\in N} e_i(g_i) + \cdots + \sum_{i_1<\cdots<i_k\in N} e_{i_1,\ldots,i_k}\left(g_{i_1},\ldots,g_{i_k}\right) + \cdots + e_{i_1,\ldots,i_n}(g_1,\ldots,g_n).$$

Substituting this expression into the definition of $V$, we obtain

$$V = \frac{1}{q^n}\sum_{\boldsymbol{g}\in G}\left[\sum_{i\in N} e_i(g_i) + \cdots + \sum_{i_1<\cdots<i_k\in N} e_{i_1,\ldots,i_k}\left(g_{i_1},\ldots,g_{i_k}\right) + \cdots + e_{i_1,i_2,\ldots,i_n}(g_1,g_2,\ldots,g_n)\right]^2. \quad \text{(A2)}$$

To simplify this equation, we prove the following lemma.

**Lemma 2**. For any two distinct site-combinations $(i_1,\ldots,i_k) \neq (j_1,\ldots,j_l)$,

$$\frac{1}{q^n}\sum_{\boldsymbol{g}\in G} e_{i_1,\ldots,i_k}\left(g_{i_1},\ldots,g_{i_k}\right) e_{j_1,\ldots,j_l}\left(g_{j_1},\ldots,g_{j_l}\right) = 0.$$

Before proving Lemma 2, let us check how it helps simplify Eq. (A2). Recall that $\left(\sum_i x_i\right)^2 = \sum_i x_i^2 + \sum_{i,j} x_i x_j$. Under Lemma 2, Eq. (A2) simplifies to

$$V = \frac{1}{q^n} \sum_{\boldsymbol{g} \in G} \left[ \sum_{i \in N} e_i(g_i)^2 + \cdots + \sum_{i_1 < \cdots < i_k \in N} e_{i_1,\ldots,i_k}\left(g_{i_1}, \ldots, g_{i_k}\right)^2 + \cdots + e_{i_1,i_2,\ldots,i_n}(g_1, g_2, \ldots, g_n)^2 \right].$$

The set $G$ can be expressed as the union $G = \bigcup_{s_1,\ldots,s_k \in Q} G_{i_1,\ldots,i_k}^{s_1,\ldots,s_k}$ for any state-combination ($s_1$, …, $s_k$) in site-combination ($i_1$, …, $i_k$). Therefore,

$$\begin{aligned}
\frac{1}{q^n} \sum_{\boldsymbol{g} \in G} e_{i_1,\ldots,i_k}\left(g_{i_1}, \ldots, g_{i_k}\right)^2 &= \frac{1}{q^n} \sum_{s_1,\ldots,s_k \in Q} \sum_{\boldsymbol{g} \in G_{i_1,\ldots,i_k}^{s_1,\ldots,s_k}} e_{i_1,\ldots,i_k}\left(g_{i_1}, \ldots, g_{i_k}\right)^2 \\
&= \frac{1}{q^n} \sum_{s_1,\ldots,s_k \in Q} \sum_{\boldsymbol{g} \in G_{i_1,\ldots,i_k}^{s_1,\ldots,s_k}} e_{i_1,\ldots,i_k}(s_1, \ldots, s_k)^2 \\
&= \frac{1}{q^n} \sum_{s_1,\ldots,s_k \in Q} q^{n-k} e_{i_1,\ldots,i_k}(s_1, \ldots, s_k)^2 \\
&= \frac{1}{q^k} \sum_{s_1,\ldots,s_k \in Q} e_{i_1,\ldots,i_k}(s_1, \ldots, s_k)^2 \\
&= Var\left(e_{i_1,\ldots,i_k} | Q^k\right).
\end{aligned}$$

This then proves the variance partition formula.

**Proof of Lemma 2.**

We make our notation flexible so that the order of sites can be permuted: for example, $e_{i_1,i_2}(s_1, s_2) = e_{i_2,i_1}(s_2, s_1)$. Assume $m$ sites are shared between $(i_1, \ldots, i_k)$ and $(j_1, \ldots, j_l)$. We can write

$$(i_1, \ldots, i_k) = (a_1, \ldots, a_m, x_1, \ldots x_{k-m}),$$
$$(j_1, \ldots, j_l) = (a_1, \ldots, a_m, y_1, \ldots y_{l-m}),$$

where $(x_1, \ldots x_{k-m}) \cap (y_1, \ldots y_{l-m}) = \emptyset$. We partition $G$ as follows:

$$G = \bigcup_{s_1,\ldots,s_m \in Q} \bigcup_{u_1,\ldots,u_{k-m} \in Q} \bigcup_{v_1,\ldots,v_{l-m} \in Q} G_{a_1,\ldots,a_m,x_1,\ldots,x_{k-m},y_1,\ldots,y_{l-m}}^{s_1,\ldots,s_m,u_1,\ldots,u_{k-m},v_1,\ldots,v_{l-m}}.$$

Then,

$$\begin{aligned}
&\frac{1}{q^n} \sum_{\boldsymbol{g} \in G} e_{i_1,\ldots,i_k}\left(g_{i_1}, \ldots, g_{i_k}\right) e_{j_1,\ldots,j_l}\left(g_{j_1}, \ldots, g_{j_l}\right) \\
&= \frac{q^{n-(k+l-m)}}{q^n} \sum_{s_1,\ldots,s_m \in Q} \sum_{u_1,\ldots,u_{k-m} \in Q} \sum_{v_1,\ldots,v_{l-m} \in Q} e_{i_1 \ldots i_k}(s_1, \ldots, s_m, u_1, \ldots, u_{k-m}) e_{j_1 \ldots j_l}(s_1, \ldots, s_m, v_1, \ldots, v_{l-m}) \\
&= \frac{1}{q^{k+l-m}} \sum_{s_1,\ldots,s_m \in Q} \sum_{u_1,\ldots,u_{k-m} \in Q} e_{i_1 \ldots i_k}(s_1, \ldots, s_m, u_1, \ldots, u_{k-m}) \sum_{v_1,\ldots,v_{l-m} \in Q} e_{j_1 \ldots j_l}(s_1, \ldots, s_m, v_1, \ldots, v_{l-m}) \\
&= 0.
\end{aligned}$$

where the last equality follows from the zero-mean property. A similar proof can be constructed for when no site is shared between $(i_1, \ldots, i_k)$ and $(j_1, \ldots, j_l)$.

*Appendix 5. Robustness to measurement noise*

Our goal is to calculate reference-free effects for an unstructured genetic architecture in which each phenotype is an independent sample from a noise distribution of zero mean and variance ω. The exact value of each effect depends on the particular instantiation of the sampling process. We are interested in the variance of each effect across the possible instantiations. Let Ω denote the set of all possible instantiations. We aim to compute

$$Var(e|\Omega),$$

which, by the definition of variance, equals

$$\langle e^2|\Omega\rangle - \langle e|\Omega\rangle^2 = \langle e^2|\Omega\rangle,$$

where the last equality follows because the expected value of any phenotype and therefore any effect is zero. We prove the following formula by mathematical induction:

$$Var(e|\Omega) = \frac{(q-1)^{O(e)}}{q^n}\omega. \quad \text{(A3)}$$

Consider the intercept:

$$\begin{aligned} Var(e_0|\Omega) &= Var\left[\frac{1}{q^n}\Sigma_{\boldsymbol{g}\in G}\, y(\boldsymbol{g})\,\middle|\Omega\right] \\ &= \frac{1}{q^{2n}}\sum_{\boldsymbol{g}\in G} Var[y(\boldsymbol{g})|\Omega] \\ &= \frac{1}{q^{2n}}\sum_{\boldsymbol{g}\in G}\omega \\ &= \frac{\omega}{q^n}. \end{aligned}$$

This is Eq. (A3) for the case when *O*(*e*) equals zero. Let us now assume that Eq. (A3) holds for effects of order (*k* – 1). Recall the definition

$$e_{i_1,\dots,i_k}(s_1,\dots,s_k) = e_{i_2,\dots,i_k}(s_2,\dots,s_k)|_{i_1}^{s_1} - e_{i_2,\dots,i_k}(s_2,\dots,s_k),$$

in which an effect of order *k* is a function of effects of order (*k* – 1). We prove that for any $j \neq i_1,\dots,i_k$,

$$e_{i_1,\dots,i_k}(s_1,\dots,s_k) = \frac{1}{q}\sum_{t\in Q} e_{i_1,\dots,i_k}(s_1,\dots,s_k)|_j^t. \quad \text{(A4)}$$

That is, an effect of order *k* for the complete space *G* is the average of the same effect calculated for the subspaces defined by conditioning on any site outside the *k* focal sites. Before proving Eq. (A4), let us see how it helps us:

$$e_{i_1,\dots,i_k}(s_1,\dots,s_k) = e_{i_2,\dots,i_k}(s_2,\dots,s_k)|_{i_1}^{s_1} - e_{i_2,\dots,i_k}(s_2,\dots,s_k)$$

$$= e_{i_2,\dots,i_k}(s_2,\dots,s_k)|_{i_1}^{s_1} - \frac{1}{q}\sum_{t\in Q} e_{i_2,\dots,i_k}(s_2,\dots,s_k)|_{i_1}^{t}$$
$$= \frac{q-1}{q} e_{i_2,\dots,i_k}(s_2,\dots,s_k)|_{i_1}^{s_1} - \frac{1}{q}\sum_{t\in Q\setminus s_1} e_{i_2,\dots,i_k}(s_2,\dots,s_k)|_{i_1}^{t}.$$

Note that the $q$ terms $e_{i_2,\dots,i_k}(s_2,\dots,s_k)|_{i_1}^{t}$, $t = 1, \dots, q$, are probabilistically independent of each other because they involve genotypes from disjoint sub-spaces. Furthermore,

$$Var\left[e_{i_2,\dots,i_k}(s_2,\dots,s_k)\middle|\Omega\right] = Var\left[\frac{1}{q}\sum_{t\in Q} e_{i_2,\dots,i_k}(s_2,\dots,s_k)|_{i_1}^{t}\,\middle|\Omega\right]$$
$$= \frac{1}{q} Var\left[e_{i_2,\dots,i_k}(s_2,\dots,s_k)|_{i_1}^{t\prime}\middle|\Omega\right],$$

where $t'$ in the last term can be any one of the $q$ states. This implies that

$$Var\left[e_{i_2,\dots,i_k}(s_2,\dots,s_k)|_{i_1}^{t\prime}\middle|\Omega\right] = q \times Var\left[e_{i_2,\dots,i_k}(s_2,\dots,s_k)\middle|\Omega\right] = \frac{q(q-1)^{k-1}}{q^n}\omega.$$

From the aforementioned probabilistic independence, it follows that

$$Var\left[e_{i_1,\dots,i_k}(s_1,\dots,s_k)\middle|\Omega\right] = \left[\left(\frac{q-1}{q}\right)^2 + \frac{q-1}{q^2}\right] \times \frac{q(q-1)^{k-1}}{q^n}\omega$$
$$= \frac{(q-1)^k}{q^n}\omega.$$

We now turn to proving Eq. (A4). We can rewrite $e_i(s)$ purely in terms of average phenotypes:

$$e_i(s) = \langle y|G_i^s\rangle - \langle y|G\rangle.$$

Similarly,

$$e_{i_1,i_2}(s_1,s_2) = \left\langle y\middle|G_{i_1,i_2}^{s_1,s_2}\right\rangle - \left\langle y\middle|G_{i_1}^{s_1}\right\rangle - \left\langle y\middle|G_{i_2}^{s_2}\right\rangle + \langle y|G\rangle.$$

In general,

$$e_{i_1,\dots,i_k}(s_1,\dots,s_k) = \left\langle y\middle|G_{i_1,\dots,i_k}^{s_1,\dots,s_k}\right\rangle - \sum_{\alpha_1<\cdots<\alpha_{k-1}\in K}\left\langle y\middle|G_{i_{\alpha_1},\dots,i_{\alpha_{k-l}}}^{s_{\alpha_1},\dots,s_{\alpha_{k-l}}}\right\rangle + \sum_{\alpha_1<\cdots<\alpha_{k-2}\in K}\left\langle y\middle|G_{i_{\alpha_1},\dots,i_{\alpha_{k-2}}}^{s_{\alpha_1},\dots,s_{\alpha_{k-2}}}\right\rangle - \cdots,$$

where $K$ is the set 1, …, $k$. More compactly,

$$e_{i_1,\dots,i_k}(s_1,\dots,s_k) = \sum_{0\le l\le k}(-1)^{k-l}\sum_{\alpha_1<\cdots<\alpha_l\in K}\left\langle y\middle|G_{i_{\alpha_1},\dots,i_{\alpha_l}}^{s_{\alpha_1},\dots,s_{\alpha_l}}\right\rangle.$$

What is important here is that $e_{i_1,\dots,i_k}(s_1,\dots,s_k)$ is a linear combination of the average phenotype of every possible subset of $G$ defined by fixing the states at one or more of the $k$ sites $i_1, \dots, i_k$. Any such average phenotype can be decomposed by fixing the state at another site $j \neq i_1,\dots,i_k$. For example,

$$\left\langle y \middle| G_{i_1}^{s_1} \right\rangle = \frac{1}{q} \sum_{t \in Q} \left\langle y \middle| G_{i_1,j}^{s_1,t} \right\rangle.$$

Therefore,

$$\begin{aligned} e_{i_1,\dots,i_k}(s_1,\dots,s_k) &= \frac{1}{q} \sum_{t \in Q} \sum_{0 \le l \le k} (-1)^{k-l} \sum_{\alpha_1 < \cdots < \alpha_l \in K} \left\langle y \middle| G_{i_{\alpha_1},\dots,i_{\alpha_l},j}^{s_{\alpha_1},\dots,s_{\alpha_l},t} \right\rangle \\ &= \frac{1}{q} \sum_{t \in Q} e_{i_1,\dots,i_k}(s_1,\dots,s_k)|_{G_j^t}. \end{aligned}$$

*Appendix 6. Unbiased estimation by regression*

We show that unbiased estimates of reference-free effects can be obtained through a two-step procedure. We first solve the optimization

$$\hat{y}_k = \operatorname{argmin} \sum_{\boldsymbol{g} \in G^*} [y(\boldsymbol{g}) - y_k(\boldsymbol{g})]^2. \tag{A5}$$

Because of the degeneracy of generalized linear decomposition, there are many solutions to this optimization. We choose any solution and enforce the zero-mean property by changing its terms without altering the predicted phenotype. We show that this normalization is always possible. We then show that $\hat{y}_k$ thus obtained is an unbiased estimate of reference-free decomposition.

We reformulate Eq. (A5) as a standard regression. Let $y$ be a vector of sampled phenotypes and $\beta$ a vector of effects to infer. We write

$$y = X\beta + \epsilon, \tag{A6}$$

where $X$ is the design matrix specifying how the effects map to phenotypes. The error $\varepsilon$ is the sum of all unmodeled higher-order effects and measurement noise. The solution to this regression is not unique: $X$ is a singular matrix because of the degeneracy of generalized linear decomposition. We can make $X$ non-singular by building in the zero-mean property. For example, we can eliminate the column of $X$ corresponding to $e_i(q)$ by coding it as

$$e_i(q) = -\sum_{t \in Q \setminus q} e_i(t).$$

Similarly, we can eliminate the column of $X$ corresponding to $e_{i_1,i_2}(s_1, q)$ by coding it as

$$e_{i_1,i_2}(s_1, q) = -\sum_{t \in Q \setminus q} e_{i_1,i_2}(s_1, t).$$

In general, every term involving state $q$ in any site can be eliminated by coding it as a linear combination of terms involving states 1 to $(q - 1)$ in accordance with the zero-mean property. The design matrix thus obtained is non-singular and can be used to infer for all terms only involving states 1 to $(q - 1)$. Estimates for terms involving state $q$ can be calculated post-hoc using the zero-mean property. This proves the existence of a solution $\hat{y}_k$ whose terms satisfy the zero-mean property.

We now show that the expected value of the error $\varepsilon$ in Eq. (A6) is zero across randomly sampled genotypes. Since the design matrix is non-singular and the errors are unbiased, by the Gauss-Markov theorem the regression estimates for terms containing states 1 to $(q - 1)$ are unbiased. The post-hoc estimates for terms containing state $q$ are also unbiased because they are linear combinations of terms containing states 1 to $(q - 1)$.

In a linear decomposition of order $k$, the error for a genotype $\boldsymbol{g}$ is given by

$$\epsilon(\boldsymbol{g}) = \sum_{i_1 < \cdots < i_{k+1} \in N} e_{i_1,\ldots,i_{k+1}}(g_{i_1}, \ldots, g_{i_{k+1}}) + \cdots + e_{1,\ldots,n}(g_1, \ldots, g_n).$$

(Measurement noise is not shown because it does not affect the expected value of $\varepsilon$.) Since genotypes are randomly sampled, it suffices to show that the expected value of $\epsilon$ is zero across all genotypes. We prove a stronger result:

$$\begin{aligned}\langle e_{i_1,\ldots,i_k} | G\rangle &= \frac{1}{q^n}\sum_{g\in G} e_{i_1,\ldots,i_k}(g_1,\ldots,g_k)\\ &= \frac{1}{q^n}\sum_{s_1,\ldots,s_k\in Q}\;\sum_{g\in G_{i_1,\ldots,i_k}^{s_1,\ldots,s_k}} e_{i_1,\ldots,i_k}(g_1,\ldots,g_k)\\ &= \frac{1}{q^n}\sum_{s_1,\ldots,s_k\in Q}\;\sum_{g\in G_{i_1,\ldots,i_k}^{s_1,\ldots,s_k}} e_{i_1,\ldots,i_k}(s_1,\ldots,s_k)\\ &= \frac{1}{q^k}\sum_{s_1,\ldots,s_k\in Q} e_{i_1,\ldots,i_k}(s_1,\ldots,s_k)\\ &= 0.\end{aligned}$$

It follows that $\langle\epsilon|G\rangle = 0$.

We now show how the zero-mean property can be enforced post hoc. Consider enforcing the zero-mean property on a first-order linear decomposition without altering the predicted phenotypes (hereafter called "normalizing"). To normalize the terms for site *i*, we first subtract from each term the mean of all terms at site *i*:

$$\delta_i(s) = e_i(s) - e_i(\cdot) \Rightarrow \delta_i(\cdot) = 0.$$

Using $\delta_i(s)$ in place of $e_i(s)$ alters the predicted phenotype of every genotype by $-e_i(\cdot)$. This can be corrected by adding $e_i(\cdot)$ to the intercept. Overall, the following modifications normalize any first-order linear decomposition:

$$\begin{aligned}\delta_i(s) &= e_i(s) - e_i(\cdot),\\ \delta_0 &= e_0 + \sum_{i\in N} e_i(\cdot).\end{aligned}$$

Similarly, the following modifications normalize any second-order linear decomposition:

$$\begin{aligned}\delta_{i_1,i_2}(s_1,s_2) &= e_{i_1,i_2}(s_1,s_2) - e_{i_1,i_2}(\cdot,s_2) - e_{i_1,i_2}(s_1,\cdot) + e_{i_1,i_2}(\cdot,\cdot),\\ \delta_i(s) &= [e_i(s) - e_i(\cdot)] + \sum_{j\in N\setminus i}\left[e_{i,j}(s,\cdot) - e_{i,j}(\cdot,\cdot)\right],\\ \delta_0 &= e_0 + \sum_{i\in N} e_i(\cdot) + \sum_{i_1<i_2\in N} e_{i_1,i_2}(\cdot,\cdot).\end{aligned}$$

For any third-order linear decomposition:

$$\begin{aligned}&\delta_{i_1,i_2,i_3}(s_1,s_2,s_3)\\ &\quad= e_{i_1,i_2,i_3}(s_1,s_2,s_3) - \left[e_{i_1,i_2,i_3}(\cdot,s_2,s_3) + e_{i_1,i_2,i_3}(s_1,\cdot,s_3) + e_{i_1,i_2,i_3}(s_1,s_2,\cdot)\right]\\ &\quad+ \left[e_{i_1,i_2,i_3}(\cdot,\cdot,s_3) + e_{i_1,i_2,i_3}(\cdot,s_2,\cdot) + e_{i_1,i_2,i_3}(s_1,\cdot,\cdot)\right] - e_{i_1,i_2,i_3}(\cdot,\cdot,\cdot),\end{aligned}$$

$$\delta_{i_1.i_2}(s_1,s_2) = e_{i_1,i_2}(s_1,s_2) - \left[e_{i_1,i_2}(\cdot,s_2) + e_{i_1,i_2}(s_1,\cdot)\right] + e_{i_1,i_2}(\cdot,\cdot)$$
$$+ \sum_{i_3 \in N \setminus i_1,i_2} \left[e_{i_1,i_2,i_3}(s_1,s_2,\cdot) - e_{i_1,i_2,i_3}(\cdot,s_2,\cdot) - e_{i_1,i_2,i_3}(s_1,\cdot,\cdot) + e_{i_1,i_2,i_3}(\cdot,\cdot,\cdot)\right],$$

$$\delta_i(s) = [e_i(s) - e_i(\cdot)] + \sum_{j \in N \setminus i} \left[e_{i,j}(s,\cdot) - \epsilon_{ij}(\cdot,\cdot)\right] + \sum_{j<k \in N \setminus i} \left[e_{i,j,k}(s,\cdot,\cdot) - \epsilon_{i,j,k}(\cdot,\cdot,\cdot)\right],$$

$$\delta_0 = \epsilon_0 + \sum_{i \in N} e_i(\cdot) + \sum_{i_1<i_2 \in N} e_{i_1,i_2}(\cdot,\cdot) + \sum_{i_1<i_2<i_3 \in N} e_{i_1,i_2,i_3}(\cdot,\cdot,\cdot).$$

Directly applying these normalization formulae is cumbersome for high orders. We provide a simple algorithm. First, we normalize the highest-order terms (order $k$) without correcting for the altered phenotypes. This can be done by Eq. (A7) shown below. Then, let $y_k$ denote the phenotype predicted by the original linear decomposition and $z_k$ the phenotypic contribution of the normalized $k$-th-order terms. We must modify the lower-order terms so that their total phenotypic contribution is $y_k - z_k$. This can be done by using regression to find a linear model of order $(k - 1)$ whose predicted phenotype is $y_k - z_k$. Such a linear model exists because post hoc enforcement of zero-mean property is always possible. Terms of order $(k - 1)$ can then be normalized and the same regression procedure can be used to correct for the altered phenotypes by changing the terms of order up to $(k - 2)$.

To show how terms of order $k$ can be normalized, we introduce a notation: $e_{i_1,\ldots,i_k}(s_1,\ldots,s_k)_{j_1,\ldots j_l}$ denotes the mean of $e_{i_1,\ldots,i_k}(s_1,\ldots,s_k)$ across sites $j_1, \ldots, j_l$. For example,

$$e_{i_1,\ldots,i_k}(s_1,\ldots,s_k)_{i_1} = e_{i_1,\ldots,i_k}(\cdot,s_2,\ldots,s_k),$$
$$e_{i_1,\ldots,i_k}(s_1,\ldots,s_k)_{i_1,i_2} = e_{i_1,\ldots,i_k}(\cdot,\cdot,s_3,\ldots,s_k).$$

Normalization of a second-order term can be restated as

$$\delta_{i_1,i_2}(s_1,s_2) = e_{i_1,i_2}(s_1,s_2) - \sum_{\alpha \in \{1,2\}} e_{i_1,i_2}(s_1,s_2)_{i_\alpha} + e_{i_1,i_2}(s_1,s_2)_{i_1,i_2}.$$

Denoting the set $\{1, \ldots, k\}$ by $K$, terms of order $k$ can be normalized by

$$\delta_{i_1,\ldots,i_k}(s_1,\ldots,s_k) = e_{i_1,\ldots,i_k}(s_1,\ldots,s_k) + \sum_{l \in K} (-1)^l \sum_{\alpha_1<\cdots<\alpha_l \in K} e_{i_1,\ldots,i_k}(s_1,\ldots,s_k)_{i_{\alpha_1},\ldots,i_{\alpha_l}}. \qquad \text{(A7)}$$
